# Obesity-induced changes in ultrastructure and calcium release of female rat cardiomyocytes are partially reversed by aerobic exercise

**DOI:** 10.64898/2026.06.18.732821

**Authors:** Anastasiia Novak, Iuliia Baglaeva, Reyhaneh Nejati Bervanlou, Bogdan Iaparov, Alexandra Zahradníková, Michal Cagalinec, Marta Novotová, Alexandra Zahradníkova

## Abstract

Obesity is associated with an elevated risk of pathological cardiac hypertrophy, whereas exercise confers cardioprotective effects; however, the cellular mechanisms underlying these opposing influences remain incompletely defined, particularly in females. We investigated how obesity and exercise affect cardiomyocyte ultrastructure, Ca²⁺ release, and contractility in female Zucker Diabetic Fatty rats and their lean littermates. Animals were assigned at 12 weeks to sedentary or aerobic exercise-trained groups and maintained on a standard diet. By 18 weeks, obese rats exhibited increased body mass and myocardial hypertrophy in the absence of diabetes. Sedentary obese animals showed a reduced fraction of compact dyads and diminished stimulated and caffeine-induced Ca²⁺ release, while contractility remained preserved. In lean rats, exercise increased dyad density but reduced Ca²⁺ release, whereas in obese rats, exercise enhanced both dyad compactness and Ca²⁺ release. Across all groups, global cardiomyocyte ultrastructure and contractile function were similar. Type III ANOVA revealed a significant obesity × exercise interaction for dyadic structure and Ca²⁺ release. These findings demonstrate that obesity itself, independent of diabetes, triggers early dyadic remodeling and altered Ca²⁺ handling in female myocardium before detectable impairment of global cardiomyocyte structure or contractile function. Furthermore, exercise exerts beneficial effects on dyadic ultrastructure and Ca²⁺ signaling in obese animals.

**New & Noteworthy:** Using a female rat model of obesity without diabetes, we demonstrate that obesity induces early remodeling of the dyadic system and impairs Ca²⁺ release in cardiac myocytes. We further show that the effects of aerobic exercise on dyadic structure and function are obesity-dependent, improving both dyad organization and Ca²⁺ signaling. These findings identify the dyadic microdomain as a vulnerable cellular site in obesity and a potential target for exercise-induced recovery.

## Introduction

Increasing evidence indicates that obesity plays an important role in the development of cardiovascular disease and heart failure (HF) (1). Obesity directly affects myocardial structure and function through a combination of hemodynamic, metabolic, and inflammatory pathways (2, 3). Nevertheless, mechanisms underlying obesity-induced cardiac pathologies at the cardiomyocyte level remain incompletely understood due to the lack of data, especially in females.

Premenopausal women are generally protected against cardiovascular disease (4–6); however, this advantage is markedly attenuated by obesity (7). In women, obesity is strongly associated with diastolic dysfunction and heart failure with preserved ejection fraction (HFpEF) (1, 8). The effects of obesity are often intermingled with the effects of diabetes; therefore, disclosing the effects of obesity in isolation remains challenging in both clinical and experimental settings. In this context, female Zucker diabetic fatty (ZDF) rats, containing a recessive mutation in the leptin receptor gene (fa), represent a valuable experimental model since, contrary to the male ZDF rats, they develop obesity but not diabetes when on a standard diet (9). This allows investigation of the obesity-associated cardiac remodeling without the influence of diabetes.

To the best of our knowledge, there are no data on the E-C coupling machinery and calcium signaling in female obese, non-diabetic rats. In obesity with diabetes, accumulating evidence indicates that cardiovascular dysfunction exhibits pronounced sexual dimorphism. Previous work in female ZDF rats, in which diabetes was induced by a high-fat diet, has demonstrated that at the age of 5 months, cardiometabolic stress can be associated with marked alterations in whole-heart structure and function, leading to early diastolic dysfunction in both sexes (8). Female diabetic ZDF rats showed significant cardiac hypertrophy and early systolic dysfunction, not observed in males (8). The study that compared male ZL (lean), ZF (obese but non-diabetic), and ZDF (obese and diabetic) rats reported that only the non-diabetic ZF group showed significant hypertrophy, while ZDF but not ZF rats showed prolonged decay of the calcium transients (10, 11). Female diabetic ZDF rats also showed impaired, while male ZDF rats showed enhanced endothelium-dependent vasorelaxation despite comparable metabolic disturbances (12). These findings indicate that biological sex modulates cardiovascular phenotypes in obesity-associated metabolic disease and underscore the importance of studying female-specific mechanisms, particularly at early disease stages (13, 14).

Exercise training is widely recognized as an effective non-pharmacological strategy to reduce cardiovascular risk and improve cardiac function in cardiometabolic disease (15, 16). Notably, the cardioprotective effects of exercise extend beyond improvements in body weight or systemic metabolic parameters. In a mouse genetic model of obesity, aerobic exercise led to improved cardiovascular function in female leptin-deficient ob/ob mice (17) and enhanced resistance to myocardial infarction in male WT and ob/ob mice independent of metabolic normalization (18). These exercise-induced cardiac adaptations have been attributed to coordinated systemic and myocardial responses, including improved metabolic flexibility, vascular function, and myocardial remodeling (19). However, how exercise influences cardiomyocyte function in the setting of obesity remains poorly defined.

Myocardial dysfunction induced by myocardial injury or overload was found to be associated with pathological changes in the structure and function of dyads of cardiomyocytes (20, 21). The dyads are specialized membrane structures that mediate excitation-contraction coupling (ECC) in cardiac muscle cells by translation of the action potential into Ca²⁺ release from the sarcoplasmic reticulum (SR) (22–24). Alterations in Ca²⁺ signals and dyadic organization have been reported in multiple forms of cardiac pathology, including cardiometabolic disease, and are thought to contribute to impaired myocardial performance (20, 25, 26). Such alterations may be particularly relevant in cardiometabolic conditions, including obesity, where subtle defects in Ca²⁺ signaling might potentially initiate cardiac dysfunction. Thus, the need for a better understanding of the effects of cardiometabolic conditions and exercise on cardiomyocyte structure-function relationships becomes evident. Therefore, we investigated the effects of obesity and exercise on cardiomyocytes by combining quantitative analysis of dyadic structure with the assessment of Ca²⁺ transients and sarcomere shortening. We have chosen the homozygous Zucker diabetic fatty rat (fa/fa) as the obese animal model, and their heterozygous lean (fa/+) littermates as controls. This allowed us to study the effects of obesity and exercise at minimal genetic variance. We focused on female rats to partially fill the gap in data on female myocardium and to eliminate diabetes as a confounding factor. This integrative approach allowed us to evaluate changes in the relationship between dyadic structure, Ca²⁺ release, and myocyte contraction due to obesity and exercise in rat female hearts.

## Methods

Zucker Diabetic Fatty (ZDF, RRID: RGD_70459, fa/fa) rat females and their lean (fa/+) littermates (*n* = 46) from Breeding Farm Dobra Voda (reg. No. SK CH 24011, Slovak Republic) were studied at 12–18 weeks of age. Animals were housed three per cage in a temperature-controlled room on a 12:12 h light/dark cycle, and provided food and water *ad libitum*. The feed mixture (Dobrá Voda, Slovakia) contained 25.5 g protein, 4.5 g fat, 5.0 g fiber, 8.1 g ash, and 54.7 g nitrogen-free extract (carbohydrates) per 100 g dry matter, providing 16 MJ/kg of metabolizable energy (327 kcal/100 g, as fed). The study protocols were reviewed approved by the Ethics Committee of Biomedical Research Center, Slovak Academy of Sciences, and by the State Veterinary and Food Administration of the Slovak Republic (protocol No: Ro 2978-5/2021-220) and conformed to the European directive 2010/63/EU and to Act No. 377/2012 of the Government of the Slovak Republic.

Randomization of animals into experimental groups (sedentary vs. running, fixation vs. isolation of myocytes, order of processing) was performed using a random number generator with uniform distribution.

All chemicals, unless stated otherwise, were obtained from Sigma-Aldrich (RRID: SCR_008988).

### Animal exercise protocol

At 11 weeks old, all animals were acclimated to the rodent treadmill system (model 47350; Ugo Basile, Gemonio, Italy), and tested 3 times (once per day) using the protocol in Figure 1A to assess compliance with treadmill running, and randomized into four experimental groups: lean sedentary (LS rats), lean exercised (LE rats), obese sedentary (OS rats), and obese exercised (OE rats). Animals averse to running were not included in the exercise groups. Starting at 12 weeks of age, animals in the exercise groups underwent training 5 days per week using a progressively increasing low-to-moderate-intensity aerobic protocol (Figure 1B). Animals in the exercise groups were processed for experiments 3 - 4 days after the final training session.

**Figure 1.**
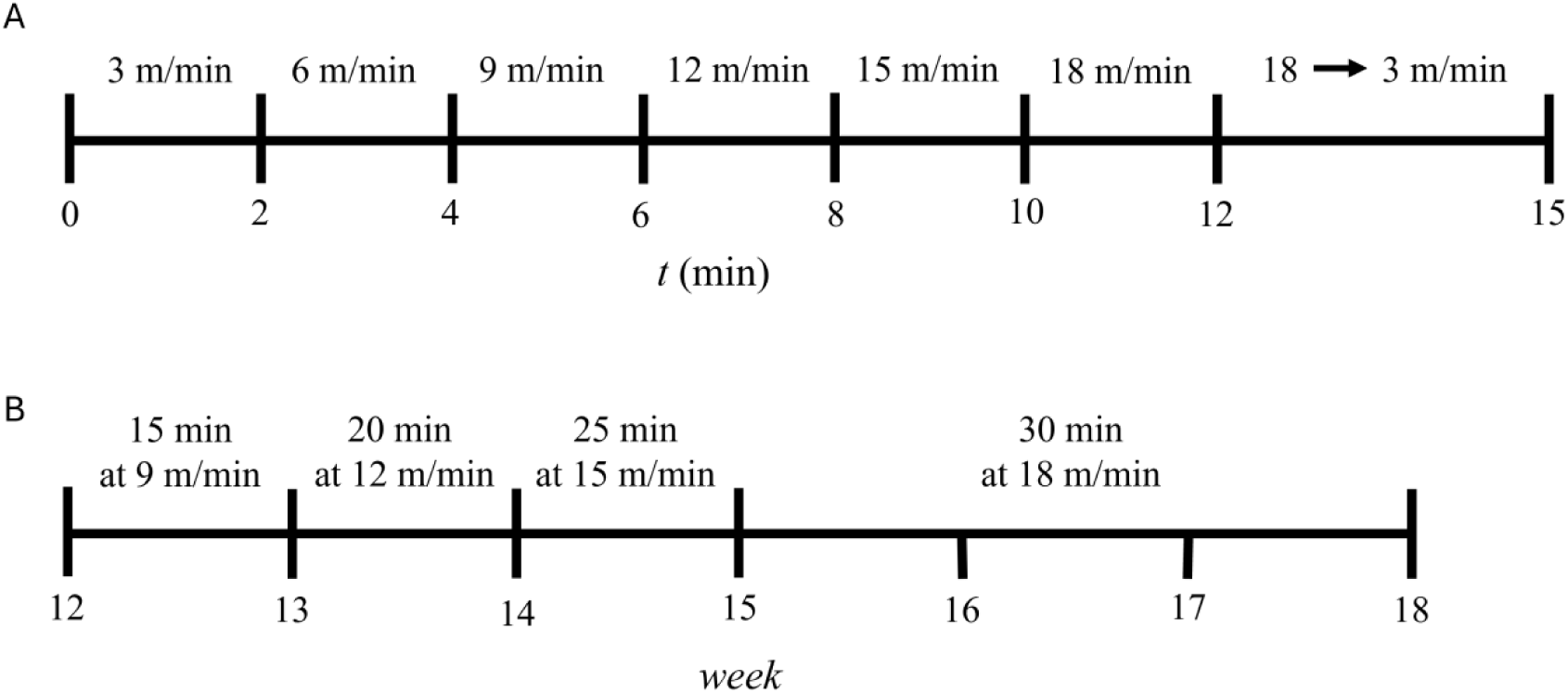
Animal exercise protocols. A – The treadmill exercise acclimation protocol; B – The experimental timeline of the training protocol (top: the duration and speed of treadmill running; bottom: the animal age axis).

At the age of 18 weeks, on the day of the experiment, the animals were weighed, the tail capillary blood was collected from 6 animals per group, and the postprandial blood glucose was measured using Accu-Chek Instant (Roche, RRID: SCR_001326) to verify the absence of diabetes. Subsequently, all animals were injected with 100 mg/kg (lean) or 150 mg/kg (obese) pentobarbital (i.p., Bioveta, a. s., Czech Republic) to induce deep anesthesia and heparin (5000 U/kg, Zentiva, a.s., Slovakia) to prevent clotting. After the loss of corneal and pedal withdrawal reflexes, the chest was opened, the heart was quickly excised, mounted by the aorta on a Langendorff-type perfusion system, and processed either for electron microscopy or isolation of cardiac myocytes (see below). After heart excision, the intraperitoneal fat weight and the right tibia length were measured. To prevent ischemic changes in ultrastructure and function, the heart weight was only measured after fixation in the hearts processed for electron microscopy.

### Electron microscopy protocol

The excised heart was perfused with air-oxygenated 37°C Tyrode solution (in mmol/l: 135 NaCl, 5.4 KCl, 1.2 MgSO_4_, 1.2 NaH_2_PO_4_, 10 HEPES, 1 CaCl_2_, pH 7.3, 300 mOsm) and left to beat for 2-3 min. Afterwards, the heart was relaxed for 5 minutes by 0Ca Tyrode solution (in mmol/l: 135 NaCl, 5.4 KCl, 1.2 MgSO_4_, 1.2 NaH_2_PO_4_, 10 HEPES; pH 7.3; 300 mOsm), supplemented with 10 mmol/l EGTA, and prefixed by 10-min perfusion with modified Karnovsky fixation buffer (2% formaldehyde, 2.5% glutaraldehyde (Electron Microscopy Sciences, USA), 150 mmol/l sodium cacodylate, 10 mmol/l EGTA; pH 7.4, 600 mOsm). The prefixed heart was demounted, and the papillary muscle was dissected and incubated in the Karnovsky buffer for 1 hour. The muscle was cut into ∼1 mm^3^ samples, postfixed in 1% OsO_4_ cacodylate buffer, and contrasted overnight in 1% uranyl acetate cacodylate buffer. The samples were dried in ethanol, permeated in propylene oxide/Durcupan ACM (1:1) mixture, mounted in Durcupan ACM, and hardened at 60°C for 72 hours. Ultrathin sections (60 nm) were cut using Power-Tome MT-XL (RMC/Sorvall, Tucson, USA) and contrasted with lead citrate before observation by transmission electron microscope JEM 1200 (Jeol, Tokyo, Japan). Images were recorded using a 2 Mpix CCD camera (Dual Vision 300 W, Gatan, Pleasanton, USA).

### Morphometric analysis

Each myocardium selected for analysis was represented by two samples dissected from the papillary muscle. From each sample, two random ultrathin sections were cut in parallel to the longitudinal axis of the muscle and observed at 30,000 × magnification. Myocyte images were captured sequentially as bands at three positions across the myocyte diameter. The band width was set to 1.2 × sarcomere length. Each myocardium was represented by 80 - 90 images covering 596 - 728 µm² of myocyte area.

The myocyte images were evaluated using a calibrated square grid comprising 154 test points separated by 250 nm (area of 0.0625 µm^2^ per point). The dyads in the image were classified according to the placement of ryanodine receptors (RyRs) and counted. Compact dyads were characterized by tight alignment of junctional sarcoplasmic reticulum cisternae with the transverse tubule, whereas in loose dyads, the cisternae were partially deflected away from the transverse tubule (20). The dyadic density (*N_A_*) was enumerated as the number of dyads of a specific class per myocyte area. The fraction of compact dyads (*f_c_*) was calculated as the ratio of the number of compact dyads to the number of all dyads in the examined myocyte. The number of analyzed image bands, myocyte profiles, and the evaluated myocyte areas for individual groups are presented in Supplementary Table 1.

### Isolation of cardiomyocytes

Cardiomyocytes were isolated as described in (27). In brief, the mounted heart was perfused with air-oxygenated 37°C solutions. First, 1Ca Tyrode solution, supplemented with (in mmol/l): 10 creatine, 10 taurine, and 10 glucose, was perfused until blood was washed out by the beating heart (usually 2-3 min). Second, 0Ca Tyrode solution, supplemented with (in mmol/l): 10 creatine, 10 taurine, 10 glucose, and 0.1 EGTA, was applied to relax the heart and weaken the extracellular matrix. Finally, the enzymatic digestion solution (0Ca Tyrode supplemented with, in mmol/l: 10 creatine, 10 taurine, 10 glucose, 0.05 CaCl_2_, and 5 mg/ml Liberase TM, Roche, RRID: SCR_001326) was perfused until the muscle became soft and light red (usually 7-10 minutes). Then, atria and the right ventricle were removed, and the left ventricle and septum were cut into small pieces, slowly triturated, gently stirred, and filtered through a 0.2 mm nylon mesh. The resulting suspension of isolated myocytes was centrifuged at 15 x g for 1 min, washed, and resuspended in 4 - 6 ml of the low calcium solution (0Ca Tyrode supplemented with, in mmol/l: 10 creatine, 10 taurine, 10 glucose, 0.05 CaCl_2_, 3 µg/ml insulin). The concentration of calcium was gradually increased from 0.05 to 0.1, 0.25, 0.5, and 1 mmol/l in 15 min intervals. All experiments were performed within 10 hours after isolation at room temperature.

### Confocal measurements

Isolated myocytes were loaded with the calcium indicator fluo-3 AM (2.5 µM, Molecular Probes, RRID: SCR_013318) for 15 min at room temperature, suspended in the experimental chamber, perfused with the perfusion solution (in mmol/l: 135 NaCl, 5.4 KCl, 1.2 MgSO_4_, 1.2 NaH_2_PO_4_, 10 HEPES, 10 Creatine, 10 Taurine, 10 glucose, 1.2 CaCl_2_, 3 µg/ml insulin; pH 7.3; 300 mOsm) at a rate of 1 ml/min, and electrically paced to contract using field stimulation by 1 Hz, 4 ms bi-phasic stimuli (MyoPacer EP, IonOptix, USA). Caffeine (20 mmol/l in the perfusion solution) was applied at the end of the stimulation protocol to measure the SR Ca^2+^ content.

The whole-cell fluorescence and transmitted light images of selected myocytes were recorded using a Zeiss LSM 510 META confocal microscope (RRID: SCR_018062) in the line-scan mode (488 nm excitation, 510 - 540 emission window, 1.9 ms/line frequency, 6000 lines, scaling 80 nm, pinhole 420 µm, and 63x oil-immersion objective). Thus, the whole-cell calcium transients and the corresponding sarcomere shortenings were recorded simultaneously.

### Analysis of fluorescence and sarcomere length recordings

Before analysis of the confocal image records, the proprietary Zeiss microscope image format and all image metadata were converted to OME-TIFF (Open Microscopy Environment Tagged Image File Format) (28) using in-house software [https://github.com/IuliiaBaglaeva/BioTIFF-Converter]. Line-scan images were transformed into calcium and contraction transients using TransientVisualizer [https://github.com/IuliiaBaglaeva/TransientVisualizer]. In brief, the background fluorescence was subtracted from the active fluorescence signal, and the spatial pixel intensities were summed to obtain the time profile of the calcium transient. The transmission light signal was analyzed using the modified Fast Fourier Transform procedure (29, 30) to estimate the sarcomere length contraction record. The temporal profiles of calcium transients and sarcomere contraction were analyzed using TransientAnalyzer (31) to estimate the amplitude and kinetic parameters. The records were included in the statistical analysis only if the measured myocyte was relaxed in the absence of electrical stimulation, the calcium transient had ΔF/F_0_ > 1 and signal-to-noise ratio SNR > 5, the sarcomere contraction was smooth, diastolic sarcomere length was > 1.6 µm, and the calcium transient preceded sarcomere shortening. The number of animals, cells, and transients is summarized in Supplementary Table 1.

### Caffeine-induced Ca²⁺ release analysis

The amplitude of the caffeine-induced Ca²⁺ release fluorescence transient (*ΔF/F_0,caf_*) was used as a measure of the SR Ca²⁺ content (23). Records of the caffeine-induced Ca²⁺ release were analyzed along the same preprocessing pipeline as electrically induced Ca²⁺ transients (see above). Recordings had to conform to the same criteria as the recordings of electrically stimulated calcium transients and contain no movement artifacts or spontaneous Ca²⁺ waves before caffeine application.

To assess the relationship between the electrically stimulated Ca²⁺ transients and caffeine-induced Ca²⁺ transients, a fractional release (FR) was calculated as:

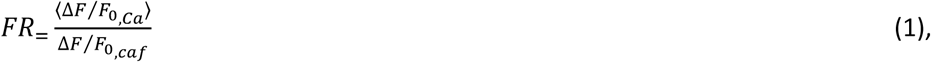

where 〈*ΔF/F_0,Ca_*〉 is the mean amplitude of electrically stimulated Ca²⁺ transients and (*ΔF/F_0,caf_*) is the amplitude of the caffeine-induced transient measured in the same cell. Experimental groups included in the analysis of caffeine-induced transients are summarized in Supplementary Table 1.

### Statistical analysis

The basic rat phenotyping parameters were compared among experimental groups using two-way Type III ANOVA.

To account for the hierarchical structure of the data (group/animal/cell/record), linear mixed-effects models (LMMs) were applied for the remaining analyses (32–34). The restricted maximum likelihood method was used for continuous variables, i.e., parameters of calcium transients, caffeine-induced Ca²⁺ transients, fractional sarcomere shortening, and the dyadic density. Degrees of freedom were approximated using Satterthwaite’s method (35). Due to the limited number of cardiomyocytes per animal (typically 1–5 cells per animal), maximum likelihood estimation (33) was used in the case of caffeine-induced Ca²⁺ transient parameters. Animal identity was included as a random intercept to account for clustering of myocytes to animals. Fractions of dyad types (counts per cell) were analyzed by a binomial generalized linear mixed model (GLMM) to account for the discrete nature of the data and the hierarchical data structure (36, 37).

Statistical significance of fixed effects (obesity, exercise) was assessed using Type III ANOVA tests.

Parameters that showed statistically significant differences in Type III ANOVA tests were compared post-hoc using Tukey’s Honestly Significant Difference (HSD) test to adjust for multiple pairwise comparisons. Statistical significance was set at p < 0.05.

Statistical analysis was implemented in R (version 4.5.0; R Core Team, 2025) using the following packages: *car* for Type III ANOVA tests; *lme4* and *lmerTest* for linear mixed-effects models (35); *glmmTMB* for fitting binomial generalized linear mixed models (38); and *emmeans* for the estimation of marginal means and Tukey’s post hoc pairwise comparisons with adjusted confidence intervals and p-values.

## Results

The animals used in this study were age-matched littermates derived from multiple litters generated by pairing fa/+ females with fa/fa males of the Zucker Diabetic Fatty rat strain, thereby ensuring a consistent genetic background across lean and obese groups.

### 1 Animal phenotyping

Animal phenotyping was performed at the end of the 18^th^ week to characterize experimental groups and to assess the effect of exercise at the organism level (Table 1). As expected, at the age of 18 weeks, the animals in the obese groups were substantially heavier (by 88%) than in the lean groups, with no difference in the tibia length. The heart weight of obese groups was about 18 % larger than that of lean groups. In the obese sedentary group, the postprandial glucose was non-significantly higher than in the remaining groups, but a significant interaction between exercise and body type (p = 0.030) was present. This was due to one of the animals having a higher blood glucose value (10 mmol/l, i.e., below the diabetic level of 13 mmol/l); disregarding this data point did not change the outcome of the analysis. The ANOVA analysis did not reveal a significant interaction between body type and exercise interventions in the remaining parameters. Overall, these data show that body type was the primary determinant of phenotypic differences, whereas the exercise intervention induced only modest effects at the organism level.

**Table 1.** Animal phenotype parameters.

| Parameter | Lean Sedentary |  | Lean Exercised |  | Obese Sedentary |  | Obese Exercised |  | P-values |  |  |
| --- | --- | --- | --- | --- | --- | --- | --- | --- | --- | --- | --- |
| | Mean ± SD | n | Mean ± SD | n | Mean ± SD | n | Mean ± SD | n | EX | BT | INT |
| BW (g) | 219 ± 17 | 12 | 215 ± 18 | 12 | 411 ± 24 | 12 | 407 ± 22 | 10 | 4.6E-01 | **2.9E-31** | 9.7E-01 |
| HW (g) | 0.95 ± 0.14 | 6 | 0.95 ± 0.17 | 4 | 1.18 ± 0.14 | 6 | 1.06 ± 0.14 | 5 | 3.7E-01 | **1.9E-02** | 3.9E-01 |
| TL (mm) | 37.3 ± 0.6 | 12 | 36.9 ± 0.9 | 12 | 37.1 ± 0.4 | 12 | 36.9 ± 0.5 | 11 | 1.2E-01 | 4.5E-01 | 6.1E-01 |
| Fat (g) | 3.5 ± 1.0 | 12 | 2.7 ± 0.6 | 12 | 33.1 ± 4.0 | 12 | 32.7 ± 4.3 | 11 | 4.9E-01 | **4.6E-33** | 8.2E-01 |
| BW/TL | 5.9 ± 0.4 | 12 | 5.8 ± 0.4 | 12 | 11.1 ± 0.6 | 12 | 11.0 ± 0.5 | 10 | 6.5E-01 | **1.9E-33** | 9.6E-01 |
| HW/TL | 0.026 ± 0.004 | 6 | 0.026 ± 0.004 | 4 | 0.032 ± 0.004 | 6 | 0.029 ± 0.004 | 5 | 4.04E-01 | **1.35E-02** | 4.43E-01 |
| Glucose PP (mM) | 5.9 ± 0.5 | 6 | 6.2 ± 0.4 | 6 | 7.1 ± 1.5 | 6 | 5.8 ± 0.3 | 6 | 1.3E-01 | 2.1E-01 | **3.0E-02** |

BW – body weight; HW – heart weight; TL – tibia length; Fat – peritoneal fat; PP – postprandial. The P-values represent the results of Type III ANOVA for the main effects of Body Type (BT), Exercise (EX), and their interaction (INT).

### 2 The effect of obesity and exercise on cardiomyocyte morphology

#### Sedentary groups

Cytoarchitecture of cardiomyocytes in lean sedentary rats appeared similar to that generally described in various rat strains kept under standard conditions (Figure 2). In ultrathin sections, myofibrils were regular and well aligned. Mitochondria were arranged in parallel with myofibrils, mostly forming single rows within the intermyofibrillar space, and exhibited an elongated shape. A network of sarcoplasmic reticulum could be observed near myofibrils. Dyads were regularly localized at the level of Z-lines.

**Figure 2.**
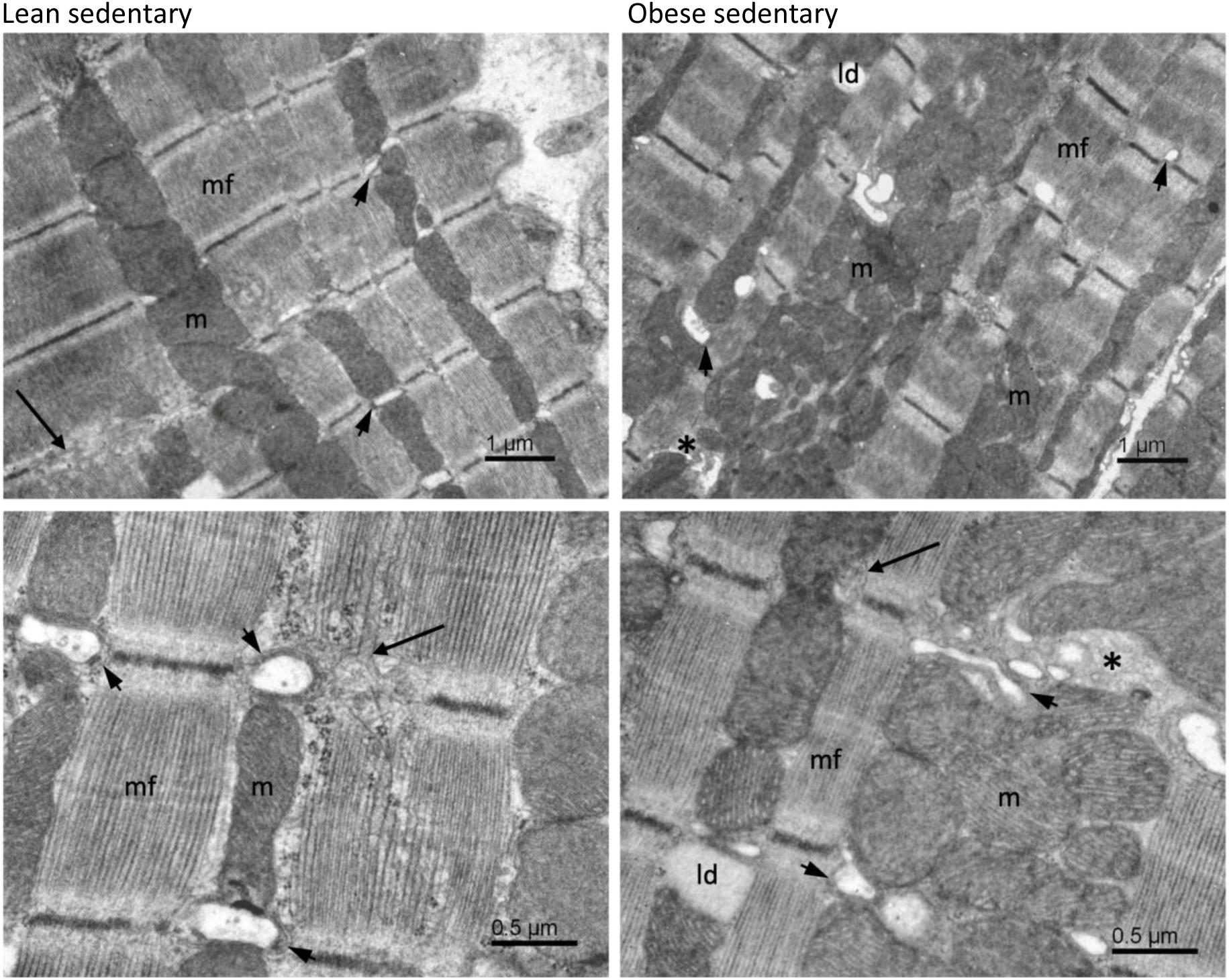
Cardiomyocyte ultrastructure in lean and obese sedentary rats. Upper row – the overview of myocyte cytoarchitecture. Lower row - details of dyads and the sarcoplasmic reticulum. mf – myofibrils; m – mitochondria; ld – lipid droplet; short arrows – dyadic microdomains; long arrows – sarcoplasmic reticulum network; asterisk – free cytoplasm.

Cardiomyocytes in obese sedentary rats were characterized by a less regular arrangement of myofibrils and dyadic microdomains than in the lean sedentary rats (Figure 2). Mitochondria often formed multiple rows or clusters. Minor constituents such as free cytoplasm and lipid droplets occurred more frequently. Altogether, the overall structural differences between myocytes of lean and obese sedentary rats were mild.

#### Exercised groups

Cardiomyocytes of exercised lean animals presented mild reorganization relative to sedentary controls. Sarcomeres of neighboring myofibrils were aligned less regularly (Figure 3). Dyads were localized near Z-lines and occasionally in duplicate. Mitochondria were also less organized and frequently formed small clusters between myofibrils. In some cells, free cytosol was occasionally present in a widened intermyofibrillar space.

**Figure 3:**
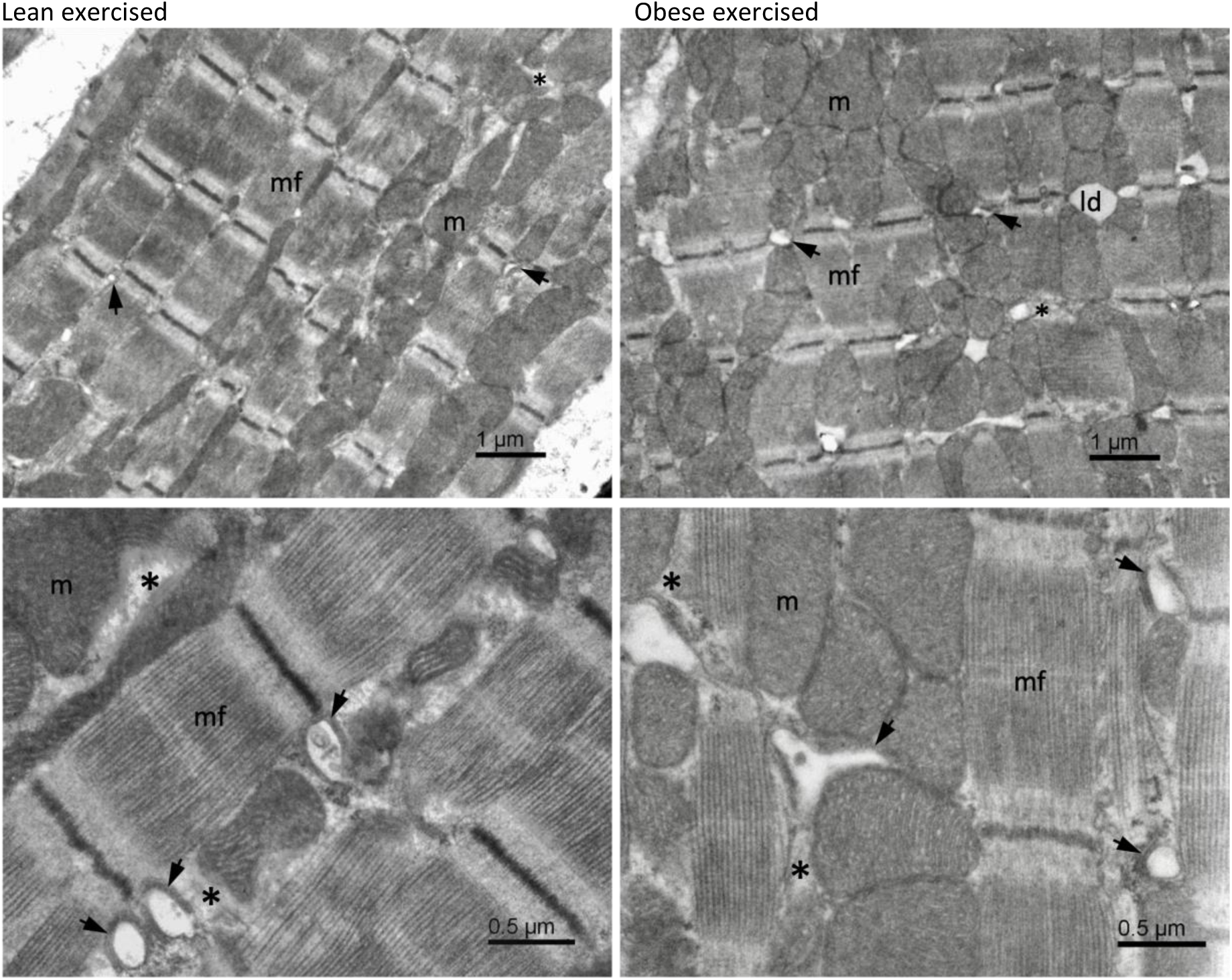
Representative electron microscopy images of cardiomyocyte ultrastructure in exercised animals. Upper row – the overview of myocyte cytoarchitecture. Lower row – details of the ultrastructure. mf – myofibrils; m – mitochondria; ld – lipid droplets; asterisks – areas of free cytoplasm; short arrows – dyads.

The morphology of cardiomyocytes in obese exercised rats (Figure 3) shared ultrastructural features with both obese sedentary and lean exercised groups. Myofibrils exhibited a slight loss of regularity in their arrangement, while overall ultrastructural organization remained preserved. Myofibrils were separated by broader bands of mitochondria. In regions with the expanded cytosolic space, clusters of mitochondria and dyadic microdomains of various sizes were observed. The presence of lipid droplets was also confirmed.

Altogether, these morphological findings indicate that the effects of obesity and exercise lead to mild adaptive remodeling of the cardiomyocyte cytoarchitecture.

### 3 The effect of obesity and exercise on dyads

Structural changes in dyads associated with obesity and exercise were quantified by classification and counting in individual myocytes in electron microscopic images of myocardial tissue samples.

Dyadic density (*N_A_*) was not different in lean and obese sedentary animals, and increased in lean but not in obese exercised animals. As a result, *N_A_* was significantly lower in obese exercised then in lean exercised animals (P=3.40×10^-5^, 4.04×10^-3^, and 8.60×10^-3^ for the effect of exercise, body type, and body type × exercise interaction, respectively, Figure 4 left, Supplementary Table 2). The largest difference in *N_A_* (Figure 4 left) was observed in lean animals, where exercise increased *N_A_* by 68% compared to sedentary conditions (p = 4.64×10^-4^). In addition, among exercised animals, *N_A_* was 34% lower in obese compared to lean animals (p = 2.21×10^-3^).

**Figure 4.**
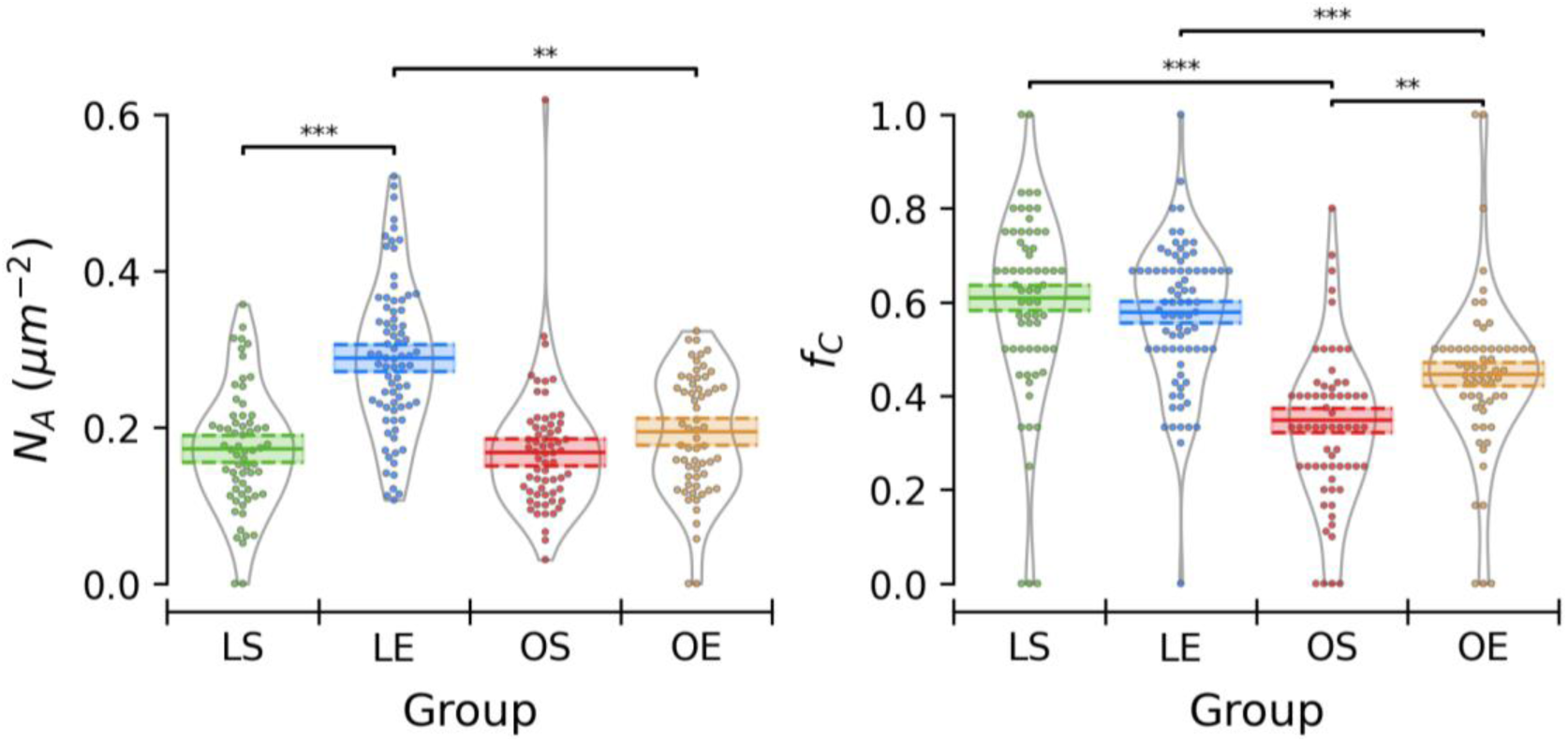
Characteristics of dyads in myocytes of experimental groups. Left – Dyadic density (*N_A_*). Right – Fraction of compact dyads (*f_C_*). LS – Lean sedentary; LE – Lean exercised; OS – Obese sedentary; OE – Obese exercised; Data points correspond to individual myocytes. Horizontal lines and bands – estimated marginal mean values and standard errors (SE) obtained from LMM for *N_A_* or GLMM for *f_C_*. Bars above plots - pairwise comparisons among groups (Tukey’s HSD test): ** p < 0.01, *** p < 0.001.

The fraction of compact dyads (*f_C_*) was significantly affected by obesity as well as by the body type × exercise interaction but not by exercise (P=1.74×10^-1^, 3.63×10^-14^, and 1.14×10^-2^ for the effect of exercise, body type, and body type × exercise interaction, respectively, Figure 4 right, Supplementary Table 2). The fraction of compact dyads was lower by 45% in obese than in lean sedentary animals (p = 1.62×10^-1^). In exercised animals, the fraction of compact dyads was less affected by obesity, and was lower by 22% in obese than in lean animals (p = 1.40×10^-4^). Therefore, even in the absence of the overall effect of exercise, it increased the fraction of compact dyads by 28% in obese animals compared to sedentary conditions (p = 7.39×10^-3^).

### 4 The effect of obesity and exercise on calcium transients and sarcomere shortening

Stimulated calcium transients and sarcomere shortenings that passed the quality criteria, recorded in 6 – 19 myocytes per animal, did not present group-specific features (Figure 5). Calcium transients and sarcomere shortenings were characterized by their amplitudes (Δ*F*/*F*_0_ or Δ*L*/*L*_0_), time to peak (*TTP*), full duration at half-maximum (*FDHM*), and the resting sarcomere length (*L_0_*).

**Figure 5.**
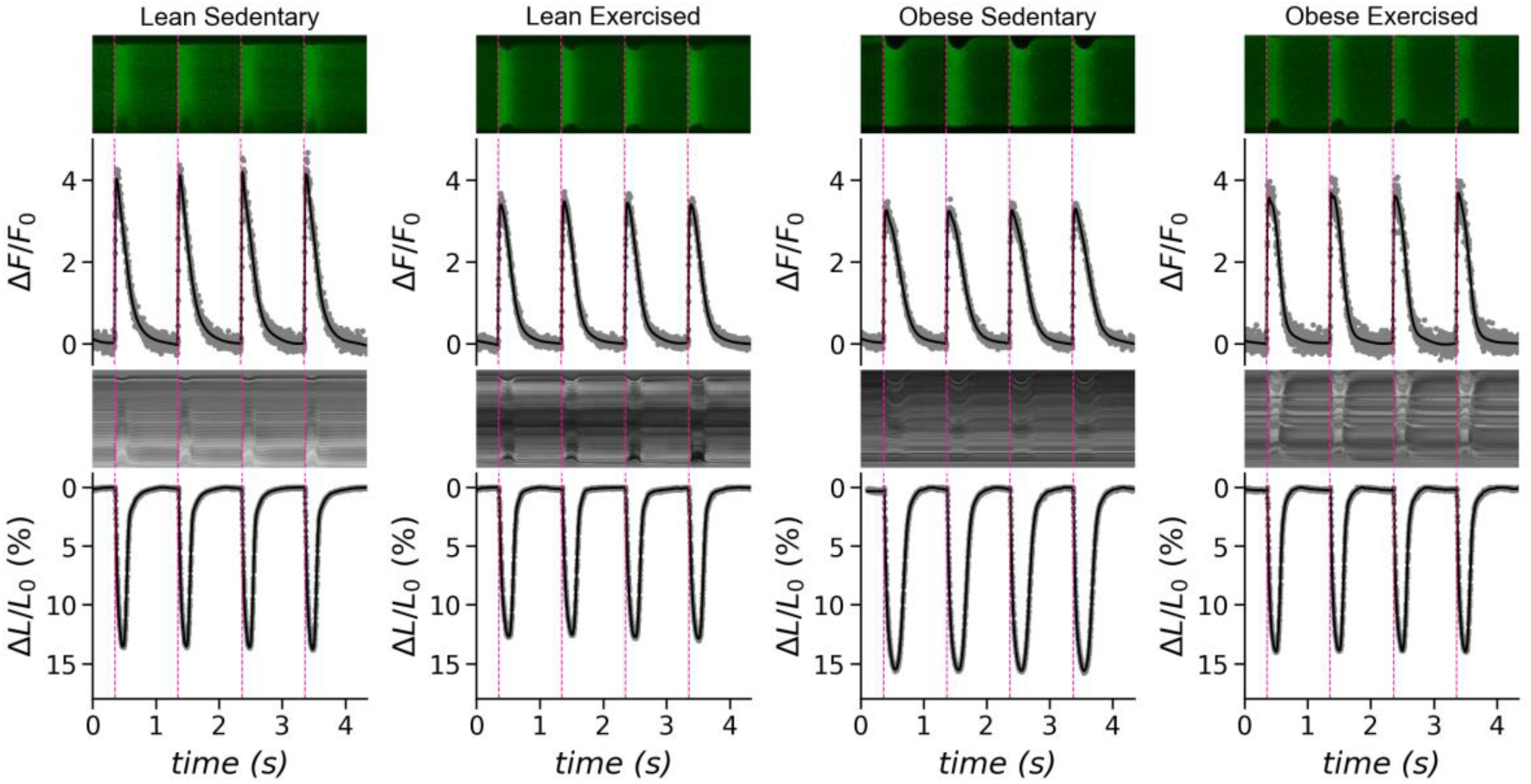
Examples of calcium transients and sarcomere shortening for individual groups. In each panel, the first row shows line-scan images of fluorescence intensity, and the traces below represent Ca²⁺ transients (ΔF/F₀). The third row shows transmission line-scan images, and the traces below represent sarcomere shortening (ΔL/L₀). Pink dashed lines correspond to the starting points of transients. Fluorescence intensity images were contrast-enhanced for clarity.

Among parameters of calcium transients, only the amplitude Δ*F*/*F*_0_ differed significantly between groups (P=1.06×10^-2^, 3.37×10^-7^, 2.86×10^-4^ for the effect of exercise, body type, and body type × exercise interaction, respectively, Figure 6 left, Supplementary Table 2). Both exercise and obesity had a significant effect on the calcium release. Interestingly, due to the obesity × exercise interaction, exercise had the opposite effect on Ca²⁺ transients in lean and obese groups. In sedentary rats, Ca²⁺ transients of the obese group were significantly lower by 24% than Ca²⁺ transients of the lean group (p = 1.5×10^-4^). Unexpectedly, the exercise reduced the amplitude Δ*F*/*F*_0_ of the lean group significantly by 13% relative to the sedentary lean group (p = 2.3×10^-2^). However, the amplitude Δ*F*/*F*_0_ of the exercised obese group was significantly higher (by 17%) than that of the sedentary obese group (p = 2.2×10^-2^). As a result, the Ca²⁺ transients of exercised obese animals reached the amplitude of exercised lean animals (p = 7.6×10^-1^). Interestingly, the changes in amplitudes were not accompanied by changes in the kinetics of Ca²⁺ transients. The differences in TTP and FDHM among the studied groups were small and did not reach statistical significance (p > 0.05) (Supplementary Table 2).

**Figure 6.**
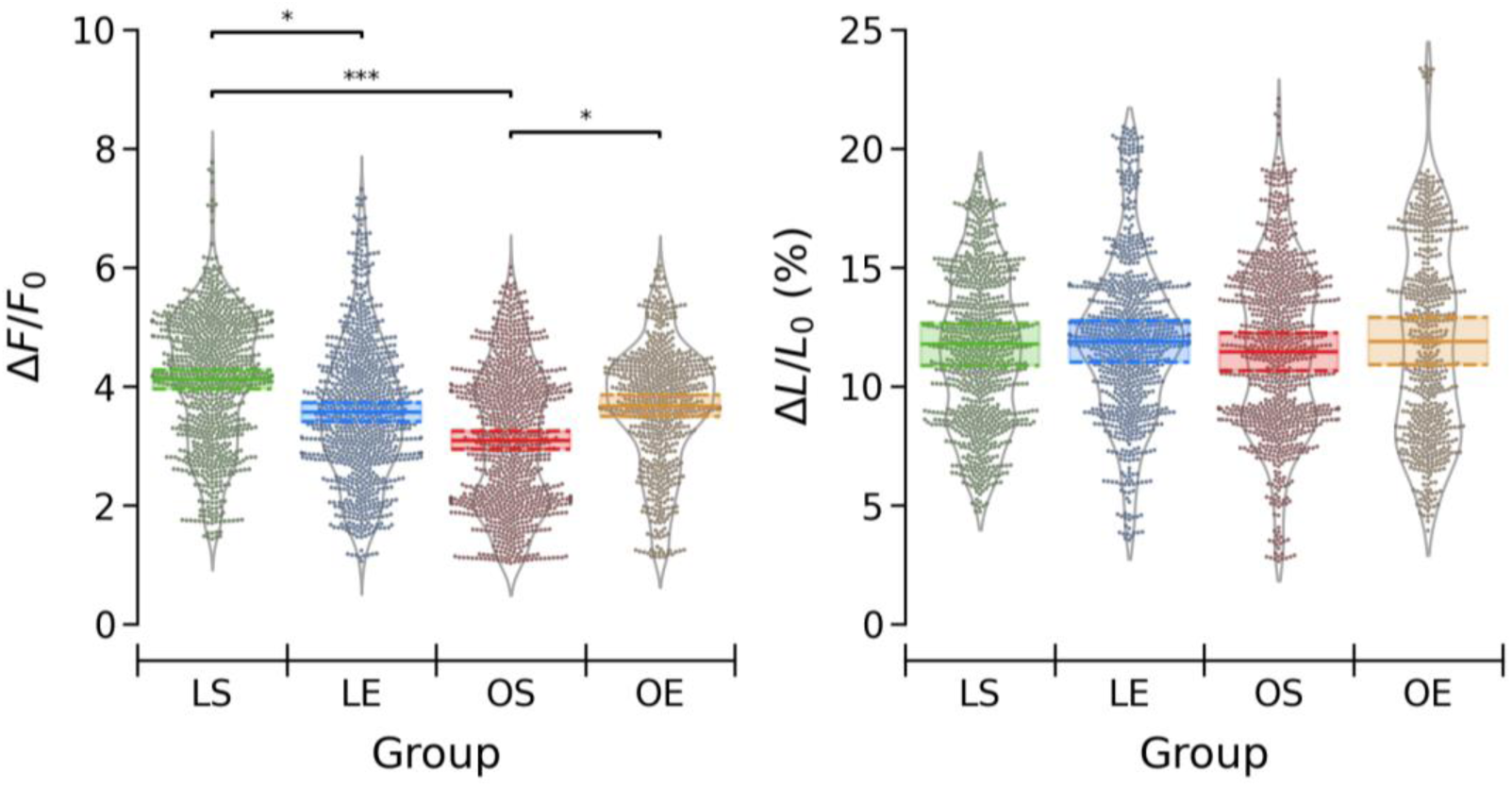
Amplitudes of calcium transients and sarcomere shortenings. Left: Calcium transients. Data points correspond to individual transients. Right: Sarcomere shortenings. Data points correspond to individual transients. LS – Lean Sedentary, LE – Lean Exercised, OS – Obese Sedentary, OE – Obese Exercised. Horizontal lines and bands – marginal mean values and standard errors (SE) estimated by LMM analysis. Bars above plots - pairwise comparisons among groups: * - p < 0.05, *** - p < 0.001.

The resting sarcomere length (L_0_), but also the amplitude of fractional shortening (ΔL/L_0_), did not differ among experimental groups (Supplementary Table 2). The sarcomere length varied by approximately 1% across all groups, while the shortening amplitude (ΔL/L_0_) differed by less than 5% (Figure 6, right). Neither the kinetic parameters of sarcomere shortenings differed significantly between the sedentary and exercised groups, both in the lean and obese rats.

TTP and FDHM of the stimulated myocyte shortenings varied substantially among individual myocytes from the same isolation but did not differ significantly among groups, as was the case with calcium transients (Supplementary Table 2).

### 5 The effect of obesity and exercise on caffeine-induced Ca²⁺ release

Caffeine-induced Ca²⁺ transients that passed the quality criteria were obtained from 1 – 5 cardiomyocytes isolated from 17 animals of either group. Representative examples of caffeine-induced Ca²⁺ transients are shown in Figure 7.

**Figure 7.**
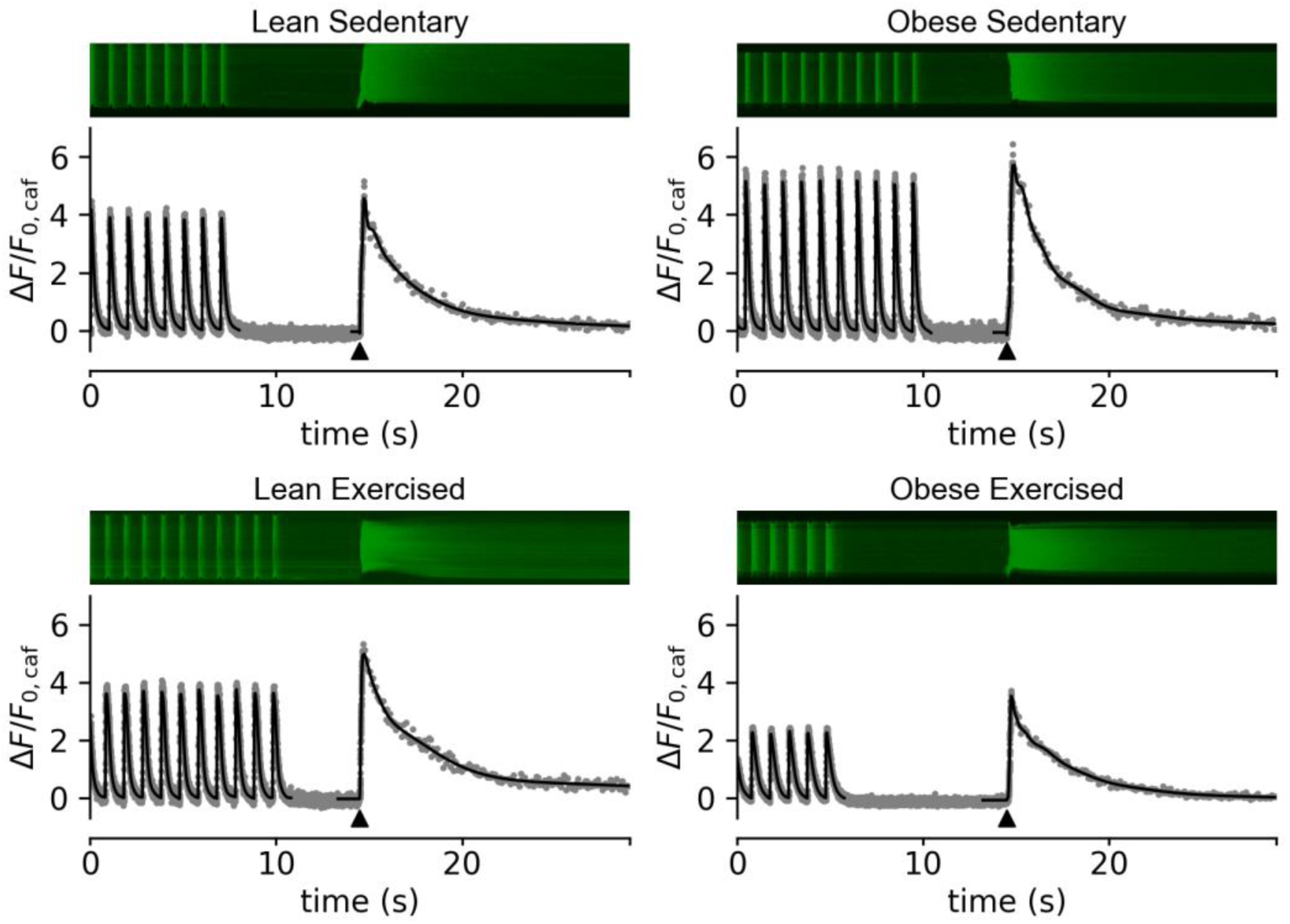
Typical records of the caffeine-induced Ca²⁺ transients. In each panel, the top rows show the line scan images and the bottom rows show the corresponding transients. Fluorescence intensity images were contrast-enhanced for clarity. Caffeine was added to the experimental chamber by perfusion at the indicated time (triangles). Only one caffeine addition was possible after the end of electrical stimulation.

Only obesity has a statistically significant effect on the caffeine-induced Ca²⁺ release amplitude (P=6.5×10^-1^, 4.7×10^-2^, 5.7×10^-1^, for the effect of exercise, body type, and body type × exercise interaction, respectively, Supplementary Table 2); however, post-hoc tests did not reveal statistically significant differences between animal groups (Figure 8, left). Both the amplitude and fractional release varied substantially between individual myocytes within groups. Nevertheless, despite the limited number of observations, it could be reasonably expected that the differences between groups would be small to marginal. Fractional release (FR) was not affected significantly (Supplementary Table 2, Figure 8, right) and varied by no more than 12% between groups.

**Figure 8.**
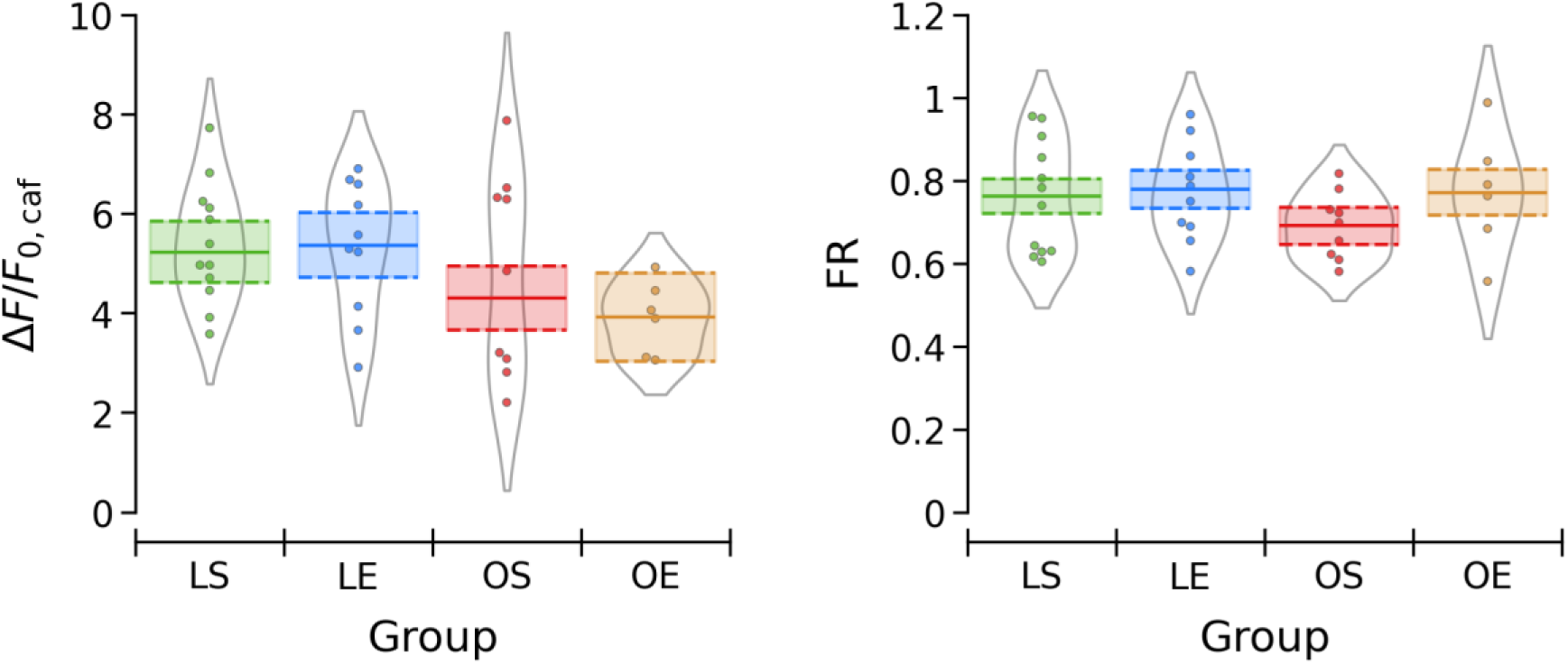
Parameters of caffeine-induced Ca²⁺ release. Left – amplitude of caffeine-induced Ca^2+^ transient (*ΔF/F_0,caf_*); Right – fractional release (FR). Data points correspond to individual cardiomyocyte transients. LS – Lean sedentary, LE – Lean exercised, OS – Obese sedentary, OE – Obese exercised. Horizontal lines and bands – the marginal mean values and standard errors (SE) estimated by LMM. No significant differences between groups were present.

## Discussion

In this study, we investigated the effects of obesity and exercise on the ultrastructure and function of the excitation-contraction coupling machinery in the female cardiac myocytes. The female Zucker Diabetic Fatty (ZDF) rat strain represents a unique model that develops obesity but not overt diabetes in the homozygous fa/fa animals (8) when maintained on a standard diet, while their heterozygous fa/+ littermates stay lean on the same diet. The major observations are summarized in Table 2.

**Table 2.** The summary of statistically highly significant effects.

| Type | Parameter | Body type<br>Obese vs. lean | Exercise<br>Lean<br>vs. sedentary | Exercise<br>Obese<br>vs. sedentary | Interaction<br>Obesity x<br>Exercise |
| --- | --- | --- | --- | --- | --- |
| Body<br>morphology | BW/TL (g) | ↑89% | - | - | - |
|  | HW/TL (g) | ↑17% | - | - | - |
| Dyads | N<sub>A</sub> | ↓22% | ↑68% | - | + |
|  | f<sub>C</sub> | ↓33% | - | ↑37% | + |
| Stimulated<br>Ca<sup>2+</sup><br>transients | ΔF/F<sub>0</sub> | ↓12% | ↓13% | ↑17% | + |
|  | kinetics | - | - | - | - |
| Contractions | L<sub>0</sub> (μm) | - | - | - | - |
|  | ΔL/L<sub>0</sub> | - | - | - | - |
| Caffeine Ca<sup>2+</sup><br>transients | ΔF/F<sub>0,caf</sub> | ↓11% | - | - | - |
|  | FR | - | - | - | - |

BW – body weight; HW – heart weight; TL – tibia length; N<sub>A</sub> – dyadic density; f<sub>C</sub> – fraction of compact dyads; ΔF/F<sub>0</sub> – calcium transient amplitude; L<sub>0</sub> (μm) – resting sarcomere length; ΔL/L<sub>0</sub> – sarcomere shortening amplitude; ΔF/F<sub>0,caf</sub> – amplitude of caffeine transient; FR – fractional release

A substantial increase in heart weight confirms the effect of obesity on cardiac hypertrophy independent of diabetes. The overall cardiomyocyte morphology in lean and obese animals kept either under sedentary or exercise regimes was not profoundly modified (Figures 2 and 3); however, substantial differences were revealed in the dyadic density and compactness of dyads (Figure 4). The obesity and exercise also affected the amplitude of calcium release (Ca^2+^ transients) induced by electrical stimulation (Figure 6, left) or application of caffeine (Figure 8). Interestingly, however, obesity and exercise had no significant effects on the resting sarcomere length, electrically stimulated sarcomere shortening (Figure 6, right), or the fractional calcium release (Figure 8). These data suggest that dyadic structures represent the element of the excitation-contraction coupling mechanism sensitive to cardiac load. Similar sensitivity of dyads was reported in the model of cardiomyocyte overload due to myocardial injury in male rats (20). The sensitivity of dyads was not fully reflected in calcium release and even less in cell contraction, probably due to calcium buffering, unloaded contractions, and eventual compensatory mechanisms.

The animal model used in this work is particularly relevant given the growing evidence that obesity alone represents an important cardiovascular risk factor in women, independent of overt diabetes (7, 39). Consistent with this study design, postprandial blood glucose levels remained below the diabetic levels across all experimental groups. At the whole-body level, obesity was associated with significant differences in multiple weight-related phenotypic parameters (Table 1), similar to the findings in female ZDF diabetic rats (8) and male ZF non-diabetic rats (10, 11).

Comparison of ultrastructural images of the sedentary lean and obese ZDF groups provided clear evidence that morphological changes induced by obesity are apparent in rat cardiac myocytes as early as at 18 weeks of age. The increased variability of myofibrillar organization, alteration in mitochondrial population, as well as widening of the free cytoplasm area, are consistent with ultrastructural changes in hypertrophied myocardium (20, 40), while the presence of lipid droplets is indicative of obesity (10, 11, 41). The exercise-induced changes in mitochondrial distribution observed in both lean and obese female ZDF rats could compensate for the morphological alterations observed in sedentary female obese ZDF animals. Comparable structural modification was also observed in diabetic female ZDF rats (8), while the increased volume fraction of lipid droplets was also observed in diabetic male ZDF rats (41). Thus, overall morphological changes appear similar in obese animals, independent of gender and diabetes.

Alterations in dyad assembly alter structural coupling between t-tubules of sarcolemma and sarcoplasmic reticulum. This compromises the quality of excitation-contraction coupling (20). In the present study, the dyad density was influenced more by exercise than by obesity, but the two factors exhibited a significant interaction. Exercise significantly increased the dyad density in lean animals but not in obese ones, indicating a reduced capacity for structural adaptation. In contrast, obesity specifically lessened the fraction of compact dyads. Although exercise did not show a main effect, it significantly, though not fully, recovered the fraction of compact dyads and counteracted the effect of obesity. This pattern suggests that obesity limits the exercise-induced increase in dyad density, while preserving the capacity of exercise to improve dyad organization, as reflected by the increased fraction of compact dyads (Figure 4).

Functionally, both obesity and exercise affected the calcium release, but with a reciprocal obesity × exercise interaction (Figure 6, left). Both factors reduced the amplitude of electrically stimulated calcium release relative to the lean sedentary group, but exercise evoked opposite responses in lean and obese animals. The effects of obesity and exercise on calcium release paralleled the effects of these modalities on the fraction of compact dyads. The amplitude of caffeine-induced Ca²⁺ release, a measure of SR Ca²⁺ content (23), was significantly reduced in obesity but not affected by other factors. This finding suggests that the obesity-associated reduction in calcium release could only partially result from its effect on the SR Ca²⁺ content.

Sarcomere shortening was not affected by obesity nor exercise (Figure 6, right) despite significant changes in Ca²⁺ release. In a previous detailed analysis comparing cardiomyocyte fractional shortening and intracellular Ca²⁺ in male diabetic ZDF and non-diabetic ZF rats (10, 11), a dissociation between Ca²⁺ release and contractile performance was also observed. In these models, Ca²⁺ transients were prolonged in diabetic (ZDF) animals without significant changes in the fractional shortening, despite marked ultrastructural remodeling (11). No significant contractility changes were observed in obese-only (ZF) rats (10, 11). Thus, both our and previous studies suggest that preserved mechanical output can coexist with impaired Ca²⁺ handling in the presence of obesity/diabesity.

Early stages of myocardial injury/heart failure have been previously observed to display very subtle changes in excitation-contraction coupling that were detectable only at the level of single dyads (20, 42). Consistent with these concepts, several experimental models of obesity and metabolic disease, including Zucker ZF and ZDF rats, exhibit reduced Ca²⁺ transient amplitude and impaired β-adrenergic responsiveness, often in the absence of overt systolic dysfunction, particularly at early disease stages (11, 43, 44).

Previous studies at the cellular, tissue, and organ levels were performed in obesity models with fully developed diabetes and in males. Some studies reported diastolic dysfunction with normal or slightly depressed systolic function and reduced developed force (45, 46), but others reported no change in single myocyte shortening (47). In field-stimulated myocytes of male diabetic rats, both calcium transients and contractions displayed a longer time to peak in young ZDF rats (48), while in older animals, only calcium transients showed prolonged time to peak (47) when compared to lean animals of the same age. In either age group, no change in the amplitude of calcium transient was observed (45, 47). Whole-heart studies revealed a decrease in conduction velocity (49), but an increase in ejection fraction and left ventricular fractional shortening (50).

Our data demonstrate that exercise and obesity influence the amplitude of Ca^2+^ release in different ways (Table 2). Obesity suppressed the amplitude of Ca^2+^ transients, suggesting impaired E-C coupling. On the other hand, exercise training restored the Ca^2+^ release amplitude in obese animals toward the lean level, resulting in a significant interaction between obesity and exercise. This pattern suggests that exercise compensated for the obesity-related disturbances in Ca²⁺ release, although full normalization was not achieved. Exercise training has previously been shown to improve cardiomyocyte Ca²⁺ handling and excitation-contraction coupling efficiency in metabolic disease models, supporting the notion that exercise can partially reverse diabetes-related impairments in Ca²⁺ signaling (51).

### Potential mechanisms underlying ECC changes

The observed alterations in Ca²⁺ release in obesity are indicative of pathological changes in excitation-contraction coupling. Importantly, this functional disorder is consistent with the observed structural remodeling of the dyadic microdomains. Such subclinical abnormalities in Ca²⁺ signaling have been proposed as early hallmarks of maladaptive cardiac remodeling (20, 42), particularly in early-stage myocardial dysfunction, where systolic function may remain preserved despite emerging defects in intracellular Ca²⁺ signaling. It may be conjectured that these changes precede the development of overt heart failure.

Alterations in Ca²⁺ transients might also arise from compromised Ca²⁺ entry through L-type calcium channels, which serve as the primary trigger for Ca²⁺ release. Electrophysiological studies in Zucker rat models have demonstrated that the L-type Ca²⁺ current density and voltage-dependent gating of L-type calcium channels are unchanged in obese (ZF) animals (52) while they are either unchanged (52) or decreased (47, 48) in diabetic obese (ZDF) animals. Early myocardial dysfunction was also associated with changes in calcium release in the absence of changes in L-type Ca^2+^ current (20, 42). In this context, the present findings suggest that the obesity-associated reduction of Ca²⁺ release is unlikely to be primarily driven by altered sarcolemmal voltage-dependent Ca²⁺ influx.

In loose dyads, the counterpart of compact dyads, a substantial fraction of RyRs is deflected from the t-tubule membrane, which delays their response to activation by L-type calcium channels (20). Considering the steep decline of the probability of RyR channel activation with increasing distance from the point Ca²⁺ source (53, 54), we can speculate that the observed decrease of Ca²⁺ transients is primarily caused by the decreased fraction of compact dyads in the myocytes of obese sedentary animals. In obese animals, exercise partially restored both the fraction of compact dyads and the Ca²⁺ transient amplitude, suggesting that exercise can mitigate obesity-associated defects in the myocyte ultrastructure-related Ca²⁺ release. Together, these findings indicate that obesity primarily drives early impairments in Ca²⁺ release that are characteristic of incipient excitation-contraction coupling dysfunction, whereas exercise acts as a modulatory factor that may delay or attenuate the progression toward overt contractile failure.

## Conclusions

On the whole, obesity intrinsically affected calcium release by dysregulation of dyads, the structural substrate of excitation-contraction coupling in cardiac myocytes. Short-term moderate intensity aerobic exercise restored both the ultrastructure of dyads and calcium release in obese female animals accustomed to a sedentary lifestyle.

## Acknowledgements

The authors thank Ladislav Novota for expert technical assistance with electron microscopy, Gizela Gajdošíková and Alžbeta Holbová for technical assistance with isolation of cardiac myocytes, Dr. Ivan Zahradník for expert insights and constructive criticism, and Dr. Stefan Zorad for help with glucose measurements.

## Author contributions

Conceived and designed research: AZ Jr, AZ

Performed experiments: IB, RNB - exercise training; RNB, MC, AZ Jr - animal phenotyping; AN, MN - electron microscopy experiments; IB, BI, MC - confocal microscopy experiments

Analyzed data: IB, BI, AN, AZ

Interpreted results of experiments: MN, AZ Jr, IB, AN, AZ

Prepared figures: MN, IB, AN, AZ

Drafted manuscript: AZ Jr, AZ

All authors edited and revised the manuscript

All authors approved the final version

## Supplementary Data

**Supplementary Table 1** contains the number of elements in the hierarchical data structures for each experimental group, as well as the areas evaluated in morphometric experiments.

**Supplementary Table 2** contains statistical analysis for each measured parameter: The raw statistics, model-estimated marginal means or probabilities, Random and Fixed effects if applicable, Type III Wald Chi-square ANOVA, and Post-hoc Tukey HSD test if applicable.

## Data Availability

Source data for this study are available from the authors upon request.

