## Supplementary Table 1 for "Obesity-induced changes in ultrastructure and calcium release of female rat cardiomyocytes are partially reversed by aerobic exercise"

[1]*Department of Cellular Cardiology, Institute of Experimental Endocrinology, Biomedical Research Center, Slovak Academy of Sciences, Bratislava, Slovakia,* [2]*Faculty of Natural Sciences, Comenius University in Bratislava, Bratislava, Slovakia,* [3]*Department of Endocrine Regulations and Neuropharmacology, Institute of Experimental Endocrinology, Biomedical Research Center, Slovak Academy of Sciences, Bratislava, Slovakia,* [4]*Laboratory of Bioinformatics, Biomedical Research Center, Slovak Academy of Sciences, Bratislava, Slovakia*

### Supplementary Table I

**Summary of experimental groups included in the morphometric analysis**

| Group | Animals | Image lines | Cell profiles | Area ($\mu m^2$) |
|---|---|---|---|---|
| LS | 4 | 11 | 64 | 2658 |
| LR | 4 | 9 | 78 | 2555 |
| OS | 4 | 12 | 64 | 2719 |
| OR | 4 | 11 | 61 | 2688 |
| Total | 16 | 43 | 267 | 10620 |

**Summary of experimental groups included in the analysis of calcium and sarcomere transients**

| Group | Animals | Cells | Calcium transients | Sarcomere shortening |
|---|---|---|---|---|
| LS | 5 | 92 | 879 | 839 |
| LR | 5 | 91 | 884 | 839 |
| OS | 6 | 103 | 972 | 940 |
| OR | 4 | 67 | 645 | 593 |
| Total | 20 | 353 | 3380 | 3211 |

**Summary of experimental groups included in the analysis of caffeine-induced calcium transients**

| Group | Animals | Cells/ Transients |
|---|---|---|
| LS | 5 | 12 |
| LR | 5 | 10 |
| OS | 5 | 9 |
| OR | 2 | 6 |
| Total | 17 | 37 |
