## Supplementary Table 2 for "Obesity-induced changes in ultrastructure and calcium release of female rat cardiomyocytes are partially reversed by aerobic exercise"

*[1]Department of Cellular Cardiology, Institute of Experimental Endocrinology, Biomedical Research Center, Slovak Academy of Sciences, Bratislava, Slovakia, [2]Faculty of Natural Sciences, Comenius University in Bratislava, Bratislava, Slovakia, [3]Department of Endocrine Regulations and Neuropharmacology, Institute of Experimental Endocrinology, Biomedical Research Center, Slovak Academy of Sciences, Bratislava, Slovakia, [4]Laboratory of Bioinformatics, Biomedical Research Center, Slovak Academy of Sciences, Bratislava, Slovakia*

#### Supplementary Table II

Significance: * - p < 0.05, ** - p < 0.01, *** - p < 0.001

#### Animal phenotyping

##### Phenotypic parameters

**Dependent Variable: BW (g)**

| Type | Factor | n | mean | SD | SE |
|---|---|---|---|---|---|
| Lean | Sedentary | 12 | 219 | 17.1 | 4.94 |
| Lean | Running | 12 | 215 | 17.8 | 5.14 |
| Obese | Sedentary | 12 | 411 | 23.6 | 6.82 |
| Obese | Running | 10 | 407 | 21.5 | 6.81 |

**Model-estimated marginal means from LM:**

| Type | Factor | Estimate | SE | df | lower.CL | upper.CL |
|---|---|---|---|---|---|---|
| Lean | Sedentary | 219.041667 | 5.81206727 | 42 | 207.31244 | 230.770893 |
| Lean | Running | 214.808333 | 5.81206727 | 42 | 203.079107 | 226.53756 |
| Obese | Sedentary | 411.308333 | 5.81206727 | 42 | 399.579107 | 423.03756 |
| Obese | Running | 406.68 | 6.3668007 | 42 | 393.831276 | 419.528724 |

**Type III two-way ANOVA:**

**Analysis of Variance Table (Type III sums of squares, F-tests)**

| Name | DF | Sum_of_Squares | Mean_Square | F_value | P_value | Sig |
|---|---|---|---|---|---|---|
| Obesity | 1 | 4.216E+05 | 4.216E+05 | 1040.08 | 2.916E-31 | *** |
| Exercise | 1 | 224.369 | 224.369 | 0.553503 | 0.461032 | |
| Interaction | 1 | 0.445786 | 0.445786 | 0.00109972 | 0.973702 | |

**Post-hoc Analysis - Tukey HSD test**

| contrast | Type/Factor | estimate | SE | df | t value | Pr(>\|t\|) | Sig |
|---|---|---|---|---|---|---|---|
| Sedentary - Running | Lean | 4.23333333 | 8.21950436 | 42 | 0.51503511 | 0.95504051 | |
| Sedentary - Running | Obese | 4.62833333 | 8.6206889 | 42 | 0.53688671 | 0.94952617 | |
| Lean - Obese | Sedentary | -192.266667 | 8.21950436 | 42 | -23.3915159 | 1.0569E-12 | *** |
| Lean - Obese | Running | -187.638333 | 8.6206889 | 42 | -21.7660486 | 1.06E-12 | *** |

### Phenotypic parameters

#### Dependent Variable: HW (g)

| Type | Factor | n | mean | SD | SE |
|---|---|---|---|---|---|
| Lean | Sedentary | 6 | 0.954 | 0.139 | 0.0569 |
| Lean | Running | 4 | 0.951 | 0.166 | 0.0829 |
| Obese | Sedentary | 6 | 1.18 | 0.14 | 0.057 |
| Obese | Running | 5 | 1.06 | 0.139 | 0.062 |

##### Model-estimated marginal means from LM:

| Type | Factor | Estimate | SE | df | lower.CL | upper.CL |
|---|---|---|---|---|---|---|
| Lean | Sedentary | 0.9535 | 0.05890889 | 17 | 0.82921311 | 1.07778689 |
| Lean | Running | 0.951 | 0.07214836 | 17 | 0.79878027 | 1.10321973 |
| Obese | Sedentary | 1.1765 | 0.05890889 | 17 | 1.05221311 | 1.30078689 |
| Obese | Running | 1.0606 | 0.06453145 | 17 | 0.92445054 | 1.19674946 |

##### Type III two-way ANOVA:

**Analysis of Variance Table (Type III sums of squares, F-tests)**

| Name | DF | Sum_of_Squares | Mean_Square | F_value | P_value | Sig |
|---|---|---|---|---|---|---|
| Obesity | 1 | 0.141221 | 0.141221 | 6.78243 | 0.0185143 | * |
| Exercise | 1 | 0.017896 | 0.017896 | 0.859496 | 0.366856 | |
| Interaction | 1 | 0.0164165 | 0.0164165 | 0.788436 | 0.386964 | |

##### Post-hoc Analysis - Tukey HSD test

| contrast | Type/Factor | estimate | SE | df | t value | Pr(>\|t\|) | Sig |
|---|---|---|---|---|---|---|---|
| Sedentary - Running | Lean | 0.0025 | 0.09314313 | 17 | 0.02684041 | 0.99999276 | |
| Sedentary - Running | Obese | 0.1159 | 0.087376 | 17 | 1.32645122 | 0.55952037 | |
| Lean - Obese | Sedentary | -0.223 | 0.08330975 | 17 | -2.67675766 | 0.06884256 | |
| Lean - Obese | Running | -0.1096 | 0.09679718 | 17 | -1.13226442 | 0.67535542 | |

### Phenotypic parameters

#### Dependent Variable: TL (mm)

| Type | Factor | n | mean | SD | SE |
|---|---|---|---|---|---|
| Lean | Sedentary | 12 | 37.3 | 0.552 | 0.159 |
| Lean | Running | 12 | 36.9 | 0.901 | 0.26 |
| Obese | Sedentary | 12 | 37.1 | 0.422 | 0.122 |
| Obese | Running | 11 | 36.9 | 0.46 | 0.139 |

##### Model-estimated marginal means from LM:

| Type | Factor | Estimate | SE | df | lower.CL | upper.CL |
|---|---|---|---|---|---|---|
| Lean | Sedentary | 37.3291667 | 0.17802358 | 43 | 36.9701479 | 37.6881854 |
| Lean | Running | 36.9483333 | 0.17802358 | 43 | 36.5893146 | 37.3073521 |
| Obese | Sedentary | 37.1 | 0.17802358 | 43 | 36.7409812 | 37.4590188 |
| Obese | Running | 36.9027273 | 0.18593957 | 43 | 36.5277444 | 37.2777102 |

##### Type III two-way ANOVA:

**Analysis of Variance Table (Type III sums of squares, F-tests)**

| Name | DF | Sum_of_Squares | Mean_Square | F_value | P_value | Sig |
|---|---|---|---|---|---|---|
| Obesity | 1 | 0.221467 | 0.221467 | 0.582334 | 0.449565 | |
| Exercise | 1 | 0.980339 | 0.980339 | 2.57775 | 0.115696 | |
| Interaction | 1 | 0.0988372 | 0.0988372 | 0.259887 | 0.612805 | |

### Phenotypic parameters

#### Dependent Variable: Fat (g)

| Type | Factor | n | mean | SD | SE |
|---|---|---|---|---|---|
| Lean | Sedentary | 12 | 3.54 | 1.03 | 0.297 |
| Lean | Running | 12 | 2.74 | 0.56 | 0.162 |
| Obese | Sedentary | 12 | 33.1 | 3.97 | 1.15 |
| Obese | Running | 11 | 32.7 | 4.28 | 1.29 |

##### Model-estimated marginal means from LM:

| Type | Factor | Estimate | SE | df | lower.CL | upper.CL |
|---|---|---|---|---|---|---|
| Lean | Sedentary | 3.54275 | 0.84907135 | 43 | 1.83043443 | 5.25506557 |
| Lean | Running | 2.73966667 | 0.84907135 | 43 | 1.02735109 | 4.45198224 |
| Obese | Sedentary | 33.1116667 | 0.84907135 | 43 | 31.3993511 | 34.8239822 |
| Obese | Running | 32.7088182 | 0.8868261 | 43 | 30.9203629 | 34.4972735 |

##### Type III two-way ANOVA:

**Analysis of Variance Table (Type III sums of squares, F-tests)**

| Name | DF | Sum_of_Squares | Mean_Square | F_value | P_value | Sig |
|---|---|---|---|---|---|---|
| Obesity | 1 | 10398 | 10398 | 1201.94 | 4.612E-33 | *** |
| Exercise | 1 | 4.26586 | 4.26586 | 0.493103 | 0.48633 | |
| Interaction | 1 | 0.469885 | 0.469885 | 0.0543152 | 0.816824 | |

##### Post-hoc Analysis - Tukey HSD test

| contrast | Type/Factor | estimate | SE | df | t value | Pr(>\|t\|) | Sig |
|---|---|---|---|---|---|---|---|
| Sedentary - Running | Lean | 0.80308333 | 1.20076822 | 43 | 0.66880795 | 0.90825453 | |
| Sedentary - Running | Obese | 0.40284848 | 1.22775515 | 43 | 0.32811794 | 0.98763467 | |
| Lean - Obese | Sedentary | -29.5689167 | 1.20076822 | 43 | -24.6249993 | 1.1001E-12 | *** |
| Lean - Obese | Running | -29.9691515 | 1.22775515 | 43 | -24.4097136 | 1.1001E-12 | *** |

### Phenotypic parameters

#### Dependent Variable: BW/TL

| Type | Factor | n | mean | SD | SE |
|---|---|---|---|---|---|
| Lean | Sedentary | 12 | 5.87 | 0.418 | 0.121 |
| Lean | Running | 12 | 5.81 | 0.383 | 0.111 |
| Obese | Sedentary | 12 | 11.1 | 0.598 | 0.173 |
| Obese | Running | 10 | 11 | 0.506 | 0.16 |

##### Model-estimated marginal means from LM:

| Type | Factor | Estimate | SE | df | lower.CL | upper.CL |
|---|---|---|---|---|---|---|
| Lean | Sedentary | 5.86572597 | 0.13931368 | 42 | 5.58457958 | 6.14687236 |
| Lean | Running | 5.80787222 | 0.13931368 | 42 | 5.52672583 | 6.08901861 |
| Obese | Sedentary | 11.0851824 | 0.13931368 | 42 | 10.804036 | 11.3663288 |
| Obese | Running | 11.0120936 | 0.15261049 | 42 | 10.7041132 | 11.3200741 |

##### Type III two-way ANOVA:

**Analysis of Variance Table (Type III sums of squares, F-tests)**

| Name | DF | Sum_of_Squares | Mean_Square | F_value | P_value | Sig |
|---|---|---|---|---|---|---|
| Obesity | 1 | 310.437 | 310.437 | 1332.92 | 1.899E-33 | *** |
| Exercise | 1 | 0.0489884 | 0.0489884 | 0.210341 | 0.648865 | |
| Interaction | 1 | 0.00066316 | 0.00066316 | 0.0028474 | 0.957697 | |

##### Post-hoc Analysis - Tukey HSD test

| contrast | Type/Factor | estimate | SE | df | t value | Pr(>\|t\|) | Sig |
|---|---|---|---|---|---|---|---|
| Sedentary - Running | Lean | 0.05785375 | 0.1970193 | 42 | 0.29364507 | 0.99105807 | |
| Sedentary - Running | Obese | 0.07308876 | 0.20663558 | 42 | 0.35370852 | 0.98461165 | |
| Lean - Obese | Sedentary | -5.21945642 | 0.1970193 | 42 | -26.4921077 | 1.0569E-12 | *** |
| Lean - Obese | Running | -5.2042214 | 0.20663558 | 42 | -25.1855047 | 1.0569E-12 | *** |

### Phenotypic parameters

#### Dependent Variable: HW/TL

| Type | Factor | n | mean | SD | SE |
|---|---|---|---|---|---|
| Lean | Sedentary | 6 | 0.026 | 0.004 | 0.00148596 |
| Lean | Running | 4 | 0.026 | 0.004 | 0.00217601 |
| Obese | Sedentary | 6 | 0.032 | 0.004 | 0.0014471 |
| Obese | Running | 5 | 0.029 | 0.004 | 0.00163069 |

##### Model-estimated marginal means from LM:

| Type | Factor | Estimate | SE | df | lower.CL | upper.CL |
|---|---|---|---|---|---|---|
| Lean | Sedentary | 0.0256376 | 0.00153095 | 17 | 0.02240758 | 0.02886762 |
| Lean | Running | 0.0255187 | 0.00187502 | 17 | 0.02156275 | 0.02947465 |
| Obese | Sedentary | 0.03151136 | 0.00153095 | 17 | 0.02828135 | 0.03474138 |
| Obese | Running | 0.02878711 | 0.00167707 | 17 | 0.0252488 | 0.03232542 |

##### Type III two-way ANOVA:

**Analysis of Variance Table (Type III sums of squares, F-tests)**

| Name | DF | Sum_of_Squares | Mean_Square | F_value | P_value | Sig |
|---|---|---|---|---|---|---|
| Obesity | 1 | 0.0001067 | 0.0001067 | 7.58717845 | 0.01354012 | * |
| Exercise | 1 | 1.0319E-05 | 1.0319E-05 | 0.73380894 | 0.40356539 | |
| Interaction | 1 | 8.6654E-06 | 8.6654E-06 | 0.61618957 | 0.44326941 | |

##### Post-hoc Analysis - Tukey HSD test

| contrast | Type/Factor | estimate | SE | df | t value | Pr(>\|t\|) | Sig |
|---|---|---|---|---|---|---|---|
| Sedentary - Running | Lean | 0.0001189 | 0.00242064 | 17 | 0.04912012 | 0.99995563 | |
| Sedentary - Running | Obese | 0.00272425 | 0.00227076 | 17 | 1.19970916 | 0.63526497 | |
| Lean - Obese | Sedentary | -0.00587376 | 0.00216509 | 17 | -2.71294531 | 0.06424786 | |
| Lean - Obese | Running | -0.00326841 | 0.0025156 | 17 | -1.29925482 | 0.57571501 | |

### Phenotypic parameters

#### Dependent Variable: Glucose PP (mM)

| Type | Factor | n | mean | SD | SE |
|---|---|---|---|---|---|
| Lean | Sedentary | 6 | 5.9 | 0.469 | 0.191 |
| Lean | Running | 6 | 6.15 | 0.373 | 0.152 |
| Obese | Sedentary | 6 | 7.1 | 1.48 | 0.604 |
| Obese | Running | 6 | 5.8 | 0.276 | 0.113 |

##### Model-estimated marginal means from LM:

| Type | Factor | Estimate | SE | df | lower.CL | upper.CL |
|---|---|---|---|---|---|---|
| Lean | Sedentary | 5.9 | 0.33084488 | 20 | 5.20986968 | 6.59013032 |
| Lean | Running | 6.15 | 0.33084488 | 20 | 5.45986968 | 6.84013032 |
| Obese | Sedentary | 7.1 | 0.33084488 | 20 | 6.40986968 | 7.79013032 |
| Obese | Running | 5.8 | 0.33084488 | 20 | 5.10986968 | 6.49013032 |

##### Type III two-way ANOVA:

**Analysis of Variance Table (Type III sums of squares, F-tests)**

| Name | DF | Sum_of_Squares | Mean_Square | F_value | P_value | Sig |
|---|---|---|---|---|---|---|
| Obesity | 1 | 1.08375 | 1.08375 | 1.65017 | 0.213616 | |
| Exercise | 1 | 1.65375 | 1.65375 | 2.51808 | 0.128232 | |
| Interaction | 1 | 3.60375 | 3.60375 | 5.48725 | 0.0296117 | * |

##### Post-hoc Analysis - Tukey HSD test

| contrast | Type/Factor | estimate | SE | df | t value | Pr(>\|t\|) | Sig |
|---|---|---|---|---|---|---|---|
| Sedentary - Running | Lean | -0.25 | 0.46788531 | 20 | -0.53431897 | 0.94964574 | |
| Sedentary - Running | Obese | 1.3 | 0.46788531 | 20 | 2.77845866 | 0.05213252 | |
| Lean - Obese | Sedentary | -1.2 | 0.46788531 | 20 | -2.56473107 | 0.07984115 | |
| Lean - Obese | Running | 0.35 | 0.46788531 | 20 | 0.74804656 | 0.87641333 | |

#### The effect of obesity and exercise on dyads

### Dyad morphometry

##### Dependent Variable: $N_A$

| Type | Factor | n | mean | SD | SE |
|---|---|---|---|---|---|
| Lean | Sedentary | 64 | 0.172 | 0.0772 | 0.00965 |
| Lean | Running | 78 | 0.289 | 0.0971 | 0.011 |
| Obese | Sedentary | 64 | 0.17 | 0.0822 | 0.0103 |
| Obese | Running | 61 | 0.191 | 0.0765 | 0.00979 |

**Model-estimated marginal means from LMM:**

| Type | Factor | Estimate | SE | df | lower.CL | upper.CL |
|---|---|---|---|---|---|---|
| Lean | Sedentary | 0.172161469 | 0.017267258 | 12.12086374 | 0.134580911 | 0.209742026 |
| Lean | Running | 0.28852436 | 0.016980367 | 11.21717635 | 0.251238917 | 0.325809804 |
| Obese | Sedentary | 0.16791448 | 0.017273575 | 12.03972454 | 0.130292362 | 0.205536598 |
| Obese | Running | 0.193984777 | 0.01741754 | 12.50512424 | 0.156204535 | 0.231765018 |

**Random effects:**

| Groups | Name | Variance | SD | Nobs |
|---|---|---|---|---|
| line_id:Animal_id | (Intercept) | 0.0002674 | 0.01635 | 43 |
| Animal_id | (Intercept) | 0.0006769 | 0.02602 | 16 |
| Residual | | 0.0064073 | 0.08005 | |

**Fixed effects:**

| Name | Estimate | SE | df | t value | Pr(>\|t\|) | Sig |
|---|---|---|---|---|---|---|
| (Intercept) | 0.205646 | 0.008591 | 11.7355 | 23.938 | 2.520E-11 | *** |
| TypeObese | 0.024697 | 0.008591 | 11.7355 | 2.875 | 0.01425 | * |
| FactorRunning | -0.035608 | 0.008591 | 11.7355 | -4.145 | 0.00142 | ** |
| TypeObese:FactorRunning | -0.022573 | 0.008591 | 11.7355 | -2.628 | 0.02242 | * |

**Type III Wald chi-square ANOVA:**

**Analysis of Deviance Table (Type III Wald chi-square tests)**

| | Chisq | df | Pr(>Chisq) | Sig |
|---|---|---|---|---|
| (Intercept) | 573.0407 | 1 | < 2.2e-16 | *** |
| Type | 8.2646 | 1 | 0.004043 | ** |
| Factor | 17.1809 | 1 | 3.398e-05 | *** |
| Type:Factor | 6.9044 | 1 | 0.008598 | *** |

**Post-hoc Analysis - Tukey HSD test**

| contrast | Type/Factor | estimate | SE | df | t value | Pr(>\|t\|) | Sig |
|---|---|---|---|---|---|---|---|
| Sedentary - Running | Lean | -0.116363 | 0.0242176 | 11.6804 | -4.80489 | 0.000463761 | *** |
| Sedentary - Running | Obese | -0.0260703 | 0.0245305 | 12.2717 | -1.06277 | 0.308354 | |
| Lean - Obese | Sedentary | 0.00424699 | 0.0244241 | 12.0808 | 0.173885 | 0.864835 | |
| Lean - Obese | Running | 0.0945396 | 0.024325 | 11.8706 | 3.88653 | 0.00220504 | ** |

### Dyad morphometry

#### Dependent Variable: $f_C$

| Type | Factor | n | mean | SD | SE |
|---|---|---|---|---|---|
| Lean | Sedentary | 64 | 0.61 | 0.0798 | 0.0399 |
| Lean | Running | 78 | 0.575 | 0.074 | 0.03684 |
| Obese | Sedentary | 64 | 0.335 | 0.043 | 0.021291 |
| Obese | Running | 61 | 0.458 | 0.0222 | 0.0111 |

##### Model-estimated probabilities from binomial GLMM:

| Type | Factor | prob | SE | df | asymp.LCL | asymp.UCL |
|---|---|---|---|---|---|---|
| Lean | Sedentary | 0.608948912 | 0.026488522 | Inf | 0.555983924 | 0.659465613 |
| Lean | Running | 0.578947569 | 0.023119066 | Inf | 0.533094082 | 0.62347846 |
| Obese | Sedentary | 0.347457253 | 0.025857764 | Inf | 0.298644425 | 0.399701584 |
| Obese | Running | 0.445933548 | 0.025667271 | Inf | 0.396343724 | 0.496623491 |

##### Random effects:

| Groups | Name | Variance | SD | Nobs |
|---|---|---|---|---|
| line_id:Animal_id | (Intercept) | 4.920E-07 | 0.0007014 | 43 |
| Animal_id | (Intercept) | 0.01262 | 0.112352 | 16 |

##### Fixed effects:

| Name | Estimate | SE | z value | Pr(>\|z\|) | Sig |
|---|---|---|---|---|---|
| (Intercept) | -0.0215 | 0.05318 | -0.404 | 0.686 | |
| TypeObese | 0.40218 | 0.0531 | 7.573 | 3.630E-14 | *** |
| FactorRunning | -0.07217 | 0.0531 | -1.359 | 0.1741 | |
| TypeObese:FactorRunning | 0.13439 | 0.05314 | 2.529 | 0.0114 | * |

##### Type III Wald chi-square ANOVA:

**Analysis of Deviance Table (Type III Wald chi-square tests)**

| | Chisq | df | Pr(>Chisq) | Sig |
|---|---|---|---|---|
| (Intercept) | 0.163465 | 1 | 0.685986 | |
| Type | 57.3571 | 1 | 3.635E-14 | *** |
| Factor | 1.84708 | 1 | 0.174123 | |
| Type:Factor | 6.3953 | 1 | 0.0114423 | * |

##### Post-hoc Analysis - Tukey HSD test

| contrast | Type/Factor | odds ratio | SE | df | z value | Pr(>\|z\|) | Sig |
|---|---|---|---|---|---|---|---|
| Sedentary - Running | Lean | 1.13252 | 0.165536 | Inf | 0.851366 | 0.394566 | |
| Sedentary - Running | Obese | 0.661583 | 0.102035 | Inf | -2.67862 | 0.00739252 | ** |
| Lean - Obese | Sedentary | 2.92452 | 0.465872 | Inf | 6.7366 | 1.621E-11 | *** |
| Lean - Obese | Running | 1.70842 | 0.240252 | Inf | 3.80841 | 0.000139865 | *** |

<u>The effect of obesity and exercise on calcium transients and sarcomere shortening</u>

#### Calcium transients

##### Dependent Variable: ΔF/$F_0$

| Type | Factor | n | mean | SD | SE |
|---|---|---|---|---|---|
| Lean | Sedentary | 879 | 4.12 | 1.12 | 0.037 |
| Lean | Running | 884 | 3.6 | 1.2 | 0.04 |
| Obese | Sedentary | 972 | 3.14 | 1.19 | 0.038 |
| Obese | Running | 645 | 3.66 | 1.03 | 0.04 |

**Model-estimated marginal means from LMM:**

| Type | Factor | Estimate | SE | df | lower.CL | upper.CL |
|---|---|---|---|---|---|---|
| Lean | Sedentary | 4.141238 | 0.15128211 | 13.8221083 | 3.81637794 | 4.46609806 |
| Lean | Running | 3.59804034 | 0.15031547 | 14.5139596 | 3.27671348 | 3.9193672 |
| Obese | Sedentary | 3.09420259 | 0.13995525 | 15.6225401 | 2.79692704 | 3.39147814 |
| Obese | Running | 3.67196733 | 0.17724231 | 14.5625533 | 3.2931935 | 4.05074117 |

**Random effects:**

| Groups | Name | Variance | SD | Nobs |
|---|---|---|---|---|
| Cell_id:Animal_id | (Intercept) | 1.26539 | 1.1249 | 353 |
| Animal_id | (Intercept) | 0.04117 | 0.2029 | 20 |
| Residual | | 0.02019 | 0.1421 | |

**Fixed effects:**

| Name | Estimate | SE | df | t value | Pr(>\|t\|) | Sig |
|---|---|---|---|---|---|---|
| (Intercept) | 4.1412 | 0.1505 | 14.2458 | 27.508 | 9.430E-14 | *** |
| TypeObese | -1.047 | 0.2052 | 15.0605 | -5.101 | 0.000129 | *** |
| FactorRunning | -0.5432 | 0.2125 | 14.5939 | -2.557 | 0.022255 | * |
| TypeObese:FactorRunning | 1.121 | 0.309 | 15.0193 | 3.628 | 0.002476 | ** |

**Type III Wald chi-square ANOVA:**

**Analysis of Deviance Table (Type III Wald chi-square tests)**

| | Chisq | df | Pr(>Chisq) | Sig |
|---|---|---|---|---|
| (Intercept) | 756.702 | 1 | < 2.2e-16 | *** |
| Type | 26.0234 | 1 | 3.373E-07 | *** |
| Factor | 6.5372 | 1 | 0.010564 | * |
| Type:Factor | 13.1608 | 1 | 0.0002859 | *** |

**Post-hoc Analysis - Tukey HSD test**

| contrast | Type/Factor | estimate | SE | df | t value | Pr(>\|t\|) | Sig |
|---|---|---|---|---|---|---|---|
| Sedentary - Running | Lean | 0.543198 | 0.213263 | 14.1603 | 2.54708 | 0.0230912 | * |
| Sedentary - Running | Obese | -0.577765 | 0.225837 | 14.9594 | -2.55833 | 0.0218689 | * |
| Lean - Obese | Sedentary | 1.04704 | 0.206092 | 14.6136 | 5.08044 | 0.000146878 | *** |
| Lean - Obese | Running | -0.073927 | 0.2324 | 14.5421 | -0.318103 | 0.754929 | |

### Calcium transients

#### Dependent Variable: TTP (ms)

| Type | Factor | n | mean | SD | SE |
|---|---|---|---|---|---|
| Lean | Sedentary | 879 | 41.2 | 12.8 | 0.43 |
| Lean | Running | 884 | 44.2 | 15 | 0.505 |
| Obese | Sedentary | 972 | 43 | 14 | 0.447 |
| Obese | Running | 645 | 41.1 | 14 | 0.552 |

##### Model-estimated marginal means from LMM:

| Type | Factor | Estimate | SE | df | lower.CL | upper.CL |
|---|---|---|---|---|---|---|
| Lean | Sedentary | 42.22917001 | 1.853381316 | 14.69990453 | 38.27174557 | 46.18659444 |
| Lean | Running | 44.31473788 | 1.833927935 | 14.8283279 | 40.40186776 | 48.227608 |
| Obese | Sedentary | 43.84879693 | 1.698788466 | 15.70762734 | 40.2420707 | 47.45552316 |
| Obese | Running | 41.2401194 | 2.151900304 | 15.69484982 | 36.6710755 | 45.8091633 |

##### Random effects:

| Groups | Name | Variance | SD | Nobs |
|---|---|---|---|---|
| Cell_id:Animal_id | (Intercept) | 110.922 | 10.532 | 353 |
| Animal_id | (Intercept) | 9.928 | 3.151 | 20 |
| Residual | | 82.805 | 9.1 | |

##### Fixed effects:

| Name | Estimate | SE | df | t value | Pr(>\|t\|) | Sig |
|---|---|---|---|---|---|---|
| (Intercept) | 42.229 | 1.847 | 13.665 | 22.86 | 2.770E-12 | *** |
| TypeObese | 1.62 | 2.508 | 14.083 | 0.646 | 0.529 | |
| FactorRunning | 2.086 | 2.601 | 13.724 | 0.802 | 0.436 | |
| TypeObese:FactorRunning | -4.694 | 3.771 | 14.172 | -1.245 | 0.233 | |

##### Type III Wald chi-square ANOVA:

**Analysis of Deviance Table (Type III Wald chi-square tests)**

| | Chisq | df | Pr(>Chisq) | Sig |
|---|---|---|---|---|
| (Intercept) | 522.598 | 1 | <2e-16 | *** |
| Type | 0.4171 | 1 | 0.5184 | |
| Factor | 0.6429 | 1 | 0.4227 | |
| Type:Factor | 1.5498 | 1 | 0.2132 | |

### Calcium transients

#### Dependent Variable: FDHM (ms)

| Type | Factor | n | mean | SD | SE |
|---|---|---|---|---|---|
| Lean | Sedentary | 879 | 201 | 19.8 | 0.669 |
| Lean | Running | 884 | 208 | 26.3 | 0.883 |
| Obese | Sedentary | 972 | 204 | 26.1 | 0.836 |
| Obese | Running | 645 | 191 | 21.7 | 0.856 |

##### Model-estimated marginal means from LMM:

| Type | Factor | Estimate | SE | df | lower.CL | upper.CL |
|---|---|---|---|---|---|---|
| Lean | Sedentary | 201.832408 | 6.00463806 | 15.7019494 | 189.083479 | 214.581338 |
| Lean | Running | 207.329645 | 5.96633234 | 15.4814006 | 194.647064 | 220.012226 |
| Obese | Sedentary | 206.325879 | 5.47014891 | 15.7598322 | 194.715303 | 217.936456 |
| Obese | Running | 190.622718 | 6.85208916 | 16.624659 | 176.14117 | 205.104266 |

##### Random effects:

| Groups | Name | Variance | SD | Nobs |
|---|---|---|---|---|
| Cell_id:Animal_id | (Intercept) | 448.19 | 21.17 | 353 |
| Animal_id | (Intercept) | 151.54 | 12.31 | 20 |
| Residual | | 15.41 | 3.925 | |

##### Fixed effects:

| Name | Estimate | SE | df | t value | Pr(>\|t\|) | Sig |
|---|---|---|---|---|---|---|
| (Intercept) | 201.832 | 6 | 16.439 | 33.64 | <2e-16 | *** |
| TypeObese | 4.493 | 8.118 | 16.467 | 0.554 | 0.587 | |
| FactorRunning | 5.497 | 8.46 | 16.324 | 0.65 | 0.525 | |
| TypeObese:FactorRunning | -21.2 | 12.176 | 16.69 | -1.741 | 0.1 | |

##### Type III Wald chi-square ANOVA:

**Analysis of Deviance Table (Type III Wald chi-square tests)**

| | Chisq | df | Pr(>Chisq) | Sig |
|---|---|---|---|---|
| (Intercept) | 1131.62 | 1 | <2e-16 | *** |
| Type | 0.3064 | 1 | 0.57992 | |
| Factor | 0.4222 | 1 | 0.51584 | |
| Type:Factor | 3.0314 | 1 | 0.08167 | |

### Sarcomere length transients

#### Dependent Variable: $L_0$ (µm)

| Type | Factor | n | mean | SD | SE |
|---|---|---|---|---|---|
| Lean | Sedentary | 839 | 1.84 | 0.0567 | 0.00196 |
| Lean | Running | 839 | 1.83 | 0.0531 | 0.00183 |
| Obese | Sedentary | 940 | 1.84 | 0.0632 | 0.00206 |
| Obese | Running | 593 | 1.86 | 0.0562 | 0.00231 |

##### Model-estimated marginal means from LMM:

| Type | Factor | Estimate | SE | df | lower.CL | upper.CL |
|---|---|---|---|---|---|---|
| Lean | Sedentary | 1.83879612 | 0.01461734 | 15.6781016 | 1.80775695 | 1.86983528 |
| Lean | Running | 1.8296511 | 0.01452044 | 15.4618938 | 1.79878185 | 1.86052034 |
| Obese | Sedentary | 1.83218196 | 0.0133107 | 15.7363855 | 1.80392607 | 1.86043784 |
| Obese | Running | 1.85062551 | 0.01669794 | 16.6273271 | 1.81533569 | 1.88591532 |

##### Random effects:

| Groups | Name | Variance | SD | Nobs |
|---|---|---|---|---|
| Cell_id:Animal_id | (Intercept) | 0.002843 | 0.053316 | 353 |
| Animal_id | (Intercept) | 0.0008869 | 0.029781 | 20 |
| Residual | | 4.867E-05 | 0.006976 | |

##### Fixed effects:

| Name | Estimate | SE | df | t value | Pr(>\|t\|) | Sig |
|---|---|---|---|---|---|---|
| (Intercept) | 1.8388 | 0.014604 | 15.6697 | 125.907 | <2e-16 | *** |
| TypeObese | -0.006614 | 0.019758 | 15.6961 | -0.335 | 0.742 | |
| FactorRunning | -0.009145 | 0.020591 | 15.5617 | -0.444 | 0.663 | |
| TypeObese:FactorRunning | 0.027589 | 0.029645 | 15.919 | 0.931 | 0.366 | |

##### Type III Wald chi-square ANOVA:

**Analysis of Deviance Table (Type III Wald chi-square tests)**

| | Chisq | df | Pr(>Chisq) | Sig |
|---|---|---|---|---|
| (Intercept) | 15852.6 | 1 | <2e-16 | *** |
| Type | 0.1121 | 1 | 0.7378 | |
| Factor | 0.1972 | 1 | 0.657 | |
| Type:Factor | 0.8661 | 1 | 0.352 | |

### Sarcomere length transients

#### Dependent Variable: $\Delta L/L_0$ (µm)

| Type | Factor | n | mean | SD | SE |
|---|---|---|---|---|---|
| Lean | Sedentary | 839 | 0.115 | 0.0323 | 0.00112 |
| Lean | Running | 839 | 0.12 | 0.0344 | 0.00119 |
| Obese | Sedentary | 940 | 0.117 | 0.0359 | 0.00117 |
| Obese | Running | 593 | 0.119 | 0.0415 | 0.00171 |

##### Model-estimated marginal means from LMM:

| Type | Factor | Estimate | SE | df | lower.CL | upper.CL |
|---|---|---|---|---|---|---|
| Lean | Sedentary | 0.1176238 | 0.00867413 | 15.6613916 | 0.09920309 | 0.13604451 |
| Lean | Running | 0.11898503 | 0.0086154 | 15.447305 | 0.10066795 | 0.13730212 |
| Obese | Sedentary | 0.11459809 | 0.00789878 | 15.7315277 | 0.09783017 | 0.13136602 |
| Obese | Running | 0.11931733 | 0.00991404 | 16.6226689 | 0.0983643 | 0.14027035 |

##### Random effects:

| Groups | Name | Variance | SD | Nobs |
|---|---|---|---|---|
| Cell_id:Animal_id | (Intercept) | 0.001035 | 0.03217 | 353 |
| Animal_id | (Intercept) | 0.0003103 | 0.017614 | 20 |
| Residual | | 1.662E-05 | 0.004076 | |

##### Fixed effects:

| Name | Estimate | SE | df | t value | Pr(>\|t\|) | Sig |
|---|---|---|---|---|---|---|
| (Intercept) | 0.117624 | 0.008666 | 15.9432 | 13.573 | 3.550E-10 | *** |
| TypeObese | -0.003026 | 0.011724 | 15.9755 | -0.258 | 0.8 | |
| FactorRunning | 0.001361 | 0.012218 | 15.8345 | 0.111 | 0.913 | |
| TypeObese:FactorRunning | 0.003358 | 0.017593 | 16.2038 | 0.191 | 0.851 | |

##### Type III Wald chi-square ANOVA:

**Analysis of Deviance Table (Type III Wald chi-square tests)**

| | Chisq | df | Pr(>Chisq) | Sig |
|---|---|---|---|---|
| (Intercept) | 184.225 | 1 | <2e-16 | *** |
| Type | 0.0666 | 1 | 0.7963 | |
| Factor | 0.0124 | 1 | 0.9113 | |
| Type:Factor | 0.0364 | 1 | 0.8486 | |

### Sarcomere length transients

#### Dependent Variable: TTP (ms)

| Type | Factor | n | mean | SD | SE |
|---|---|---|---|---|---|
| Lean | Sedentary | 839 | 142 | 23.3 | 0.803 |
| Lean | Running | 839 | 154 | 32.9 | 1.13 |
| Obese | Sedentary | 940 | 137 | 24.5 | 0.799 |
| Obese | Running | 593 | 133 | 29 | 1.19 |

##### Model-estimated marginal means from LMM:

| Type | Factor | Estimate | SE | df | lower.CL | upper.CL |
|---|---|---|---|---|---|---|
| Lean | Sedentary | 144.874706 | 8.38378233 | 15.8814797 | 127.091095 | 162.658316 |
| Lean | Running | 154.012286 | 8.34939143 | 15.6884877 | 136.283761 | 171.740812 |
| Obese | Sedentary | 137.952094 | 7.63851549 | 15.8299578 | 121.745016 | 154.159171 |
| Obese | Running | 136.524471 | 9.49373595 | 16.5501662 | 116.452887 | 156.596055 |

##### Random effects:

| Groups | Name | Variance | SD | Nobs |
|---|---|---|---|---|
| Cell_id:Animal_id | (Intercept) | 483.34 | 21.985 | 353 |
| Animal_id | (Intercept) | 319.62 | 17.878 | 20 |
| Residual | | 53.61 | 7.322 | |

##### Fixed effects:

| Name | Estimate | SE | df | t value | Pr(>\|t\|) | Sig |
|---|---|---|---|---|---|---|
| (Intercept) | 144.875 | 8.381 | 15.951 | 17.285 | 9.420E-12 | *** |
| TypeObese | -6.923 | 11.339 | 15.928 | -0.61 | 0.55 | |
| FactorRunning | 9.138 | 11.83 | 15.854 | 0.772 | 0.451 | |
| TypeObese:FactorRunning | -10.565 | 16.979 | 16.098 | -0.622 | 0.542 | |

##### Type III Wald chi-square ANOVA:

**Analysis of Deviance Table (Type III Wald chi-square tests)**

| | Chisq | df | Pr(>Chisq) | Sig |
|---|---|---|---|---|
| (Intercept) | 298.788 | 1 | <2e-16 | *** |
| Type | 0.3727 | 1 | 0.5415 | |
| Factor | 0.5966 | 1 | 0.4399 | |
| Type:Factor | 0.3872 | 1 | 0.5338 | |

### Sarcomere length transients

#### Dependent Variable: FDHM (ms)

| Type | Factor | n | mean | SD | SE |
|---|---|---|---|---|---|
| Lean | Sedentary | 839 | 190 | 37.8 | 1.31 |
| Lean | Running | 839 | 212 | 58.8 | 2.03 |
| Obese | Sedentary | 940 | 185 | 42.8 | 1.4 |
| Obese | Running | 593 | 173 | 43.9 | 1.8 |

##### Model-estimated marginal means from LMM:

| Type | Factor | Estimate | SE | df | lower.CL | upper.CL |
|---|---|---|---|---|---|---|
| Lean | Sedentary | 196.071238 | 14.8266523 | 15.8858172 | 164.621768 | 227.520707 |
| Lean | Running | 210.567583 | 14.7675907 | 15.6968588 | 179.212481 | 241.922685 |
| Obese | Sedentary | 188.178995 | 13.5088156 | 15.8317578 | 159.516832 | 216.841159 |
| Obese | Running | 178.556462 | 16.7853701 | 16.5448696 | 143.068071 | 214.044854 |

##### Random effects:

| Groups | Name | Variance | SD | Nobs |
|---|---|---|---|---|
| Cell_id:Animal_id | (Intercept) | 1488.9 | 38.586 | 353 |
| Animal_id | (Intercept) | 1002.28 | 31.659 | 20 |
| Residual | | 14.77 | 3.843 | |

##### Fixed effects:

| Name | Estimate | SE | df | t value | Pr(>\|t\|) | Sig |
|---|---|---|---|---|---|---|
| (Intercept) | 196.071 | 14.822 | 15.839 | 13.228 | 5.640E-10 | *** |
| TypeObese | -7.892 | 20.054 | 15.815 | -0.394 | 0.699 | |
| FactorRunning | 14.496 | 20.922 | 15.745 | 0.693 | 0.498 | |
| TypeObese:FactorRunning | -24.119 | 30.026 | 15.982 | -0.803 | 0.434 | |

##### Type III Wald chi-square ANOVA:

**Analysis of Deviance Table (Type III Wald chi-square tests)**

| | Chisq | df | Pr(>Chisq) | Sig |
|---|---|---|---|---|
| (Intercept) | 174.98 | 1 | <2e-16 | *** |
| Type | 0.1549 | 1 | 0.6939 | |
| Factor | 0.4801 | 1 | 0.4884 | |
| Type:Factor | 0.6452 | 1 | 0.4218 | |

<u>The effect of obesity and exercise on caffeine-induced $Ca^{2+}$ release</u>

#### Caffeine transients

##### Dependent Variable: $\Delta F/F_{0,\,caf}$

| Type | Factor | n | mean | SD | SE |
|---|---|---|---|---|---|
| Lean | Sedentary | 12 | 5.39 | 1.21 | 0.35 |
| Lean | Running | 9 | 4.79 | 2.03 | 0.678 |
| Obese | Sedentary | 10 | 5.31 | 1.36 | 0.431 |
| Obese | Running | 6 | 3.91 | 0.734 | 0.3 |

**Model-estimated marginal means from LMM:**

| Type | Factor | Estimate | SE | df | lower.CL | upper.CL |
|---|---|---|---|---|---|---|
| Lean | Sedentary | 5.29352793 | 0.61517618 | 26.2435618 | 4.02958611 | 6.55746974 |
| Lean | Running | 5.35627692 | 0.64445111 | 28.1991935 | 4.03659863 | 6.6759552 |
| Obese | Sedentary | 4.49276293 | 0.64224142 | 28.0708586 | 3.17734061 | 5.80818525 |
| Obese | Running | 3.91379736 | 0.88871434 | 23.4934392 | 2.07748594 | 5.75010877 |

**Random effects:**

| Groups | Name | Variance | SD | Nobs |
|---|---|---|---|---|
| Animal_id | (Intercept) | 0.4862 | 0.6973 | 17 |
| Residual | | 1.4443 | 1.2018 | |

**Fixed effects:**

| Name | Estimate | SE | df | t value | Pr(>\|t\|) | Sig |
|---|---|---|---|---|---|---|
| (Intercept) | 4.7641 | 0.2819 | 7.6712 | 16.902 | 2.400E-07 | *** |
| TypeObese | 0.5608 | 0.2819 | 7.6712 | 1.99 | 0.0834 | |
| FactorRunning | 0.1291 | 0.2819 | 7.6712 | 0.458 | 0.6597 | |
| TypeObese:FactorRunning | -0.1604 | 0.2819 | 7.6712 | -0.569 | 0.5855 | |

**Type III Wald chi-square ANOVA:**

**Analysis of Deviance Table (Type III Wald chi-square tests)**

| | Chisq | df | Pr(>Chisq) | Sig |
|---|---|---|---|---|
| (Intercept) | 285.667 | 1 | <2e-16 | *** |
| Type | 3.9585 | 1 | 0.04663 | * |
| Factor | 0.2096 | 1 | 0.64706 | |
| Type:Factor | 0.3239 | 1 | 0.56925 | |

**Post-hoc Analysis - Tukey HSD test**

| contrast | Type/Factor | estimate | SE | df | t value | Pr(>\|t\|) | Sig |
|---|---|---|---|---|---|---|---|
| Sedentary - Running | Lean | -0.06274899 | 0.89093152 | 27.2835288 | -0.07043077 | 0.99999947 | |
| Sedentary - Running | Obese | 0.57896557 | 1.09648859 | 25.1301248 | 0.52801787 | 0.99002835 | |
| Lean - Obese | Sedentary | 0.80076499 | 0.88933446 | 27.2323124 | 0.90040928 | 0.90523874 | |
| Lean - Obese | Running | 1.44247956 | 1.09778432 | 25.1408322 | 1.31399176 | 0.67377754 | |

### Caffeine transients

#### Dependent Variable: FR

| Type | Factor | n | mean | SD | SE |
|---|---|---|---|---|---|
| Lean | Sedentary | 12 | 0.759 | 0.135 | 0.039 |
| Lean | Running | 9 | 0.69 | 0.0801 | 0.0267 |
| Obese | Sedentary | 10 | 0.77 | 0.12 | 0.0379 |
| Obese | Running | 6 | 0.771 | 0.146 | 0.0596 |

##### Model-estimated marginal means from LMM:

| Type | Factor | Estimate | SE | df | lower.CL | upper.CL |
|---|---|---|---|---|---|---|
| Lean | Sedentary | 0.7592141 | 0.04128199 | 13.0992613 | 0.67009844 | 0.84832976 |
| Lean | Running | 0.77042419 | 0.04597222 | 14.9229335 | 0.67239264 | 0.86845575 |
| Obese | Sedentary | 0.68983164 | 0.0447565 | 19.1033829 | 0.59618949 | 0.78347378 |
| Obese | Running | 0.7708681 | 0.05445043 | 12.6273412 | 0.65288122 | 0.88885497 |

##### Random effects:

| Groups | Name | Variance | SD | Nobs |
|---|---|---|---|---|
| Animal_id | (Intercept) | 0 | 0 | 17 |
| Residual | | 0.0132 | 0.1149 | |

##### Fixed effects:

| Name | Estimate | SE | df | t value | Pr(>\|t\|) | Sig |
|---|---|---|---|---|---|---|
| (Intercept) | 0.74758 | 0.01951 | 37 | 38.327 | <2e-16 | *** |
| TypeObese | 0.01723 | 0.01951 | 37 | 0.884 | 0.383 | |
| FactorRunning | -0.02306 | 0.01951 | 37 | -1.182 | 0.245 | |
| TypeObese:FactorRunning | 0.01746 | 0.01951 | 37 | 0.895 | 0.377 | |

##### Type III Wald chi-square ANOVA:

**Analysis of Deviance Table (Type III Wald chi-square tests)**

| | Chisq | df | Pr(>Chisq) | Sig |
|---|---|---|---|---|
| (Intercept) | 1468.99 | 1 | <2e-16 | *** |
| Type | 0.7807 | 1 | 0.3769 | |
| Factor | 1.3979 | 1 | 0.2371 | |
| Type:Factor | 0.801 | 1 | 0.3708 | |
